# Alzheimer’s disease risk factor APOE4 exerts dimorphic effects on female bone

**DOI:** 10.1101/2025.10.16.682922

**Authors:** Charles A. Schurman, Gurcharan Kaur, Serra Kaya, Joanna Bons, Carlos Galicia Aguirre, Qi Liu, Christina D. King, Kenneth A. Wilson, Harrison L. Baker, Mikayala Hady, Nadja Maldonado Luna, Gregor Bieri, Saul A. Villeda, Lisa M. Ellerby, Birgit Schilling, Tamara Alliston

## Abstract

Individuals diagnosed with Alzheimer’s disease (AD) are at an increased risk of bone fractures. Conversely, a diagnosis of osteoporosis in women is the earliest known predictor for AD. However, mechanisms responsible for the coupled decline in cognitive and skeletal health remain unclear. Proteomic analysis of cortical bone from aged mice revealed neurological disease-associated proteins that are highly enriched in aged mouse bones, including apolipoprotein E (Apoe) and amyloid precursor protein. Further, Apoe localized specifically to bone-embedded osteocytes with expression twice as high in aged female bone as in young or male counterparts. In humans, *APOE* allele variants carry differing AD risk with age. To investigate *APOE* allelic roles in bone, we utilized a humanized APOE knock-in mouse model that expresses either the protective APOE2, the neutral APOE3, or the AD risk factor APOE4, and analyzed bone and hippocampus from the same mice. APOE4 exerted strong sex-specific effects on the bone transcriptome and proteome, relative to APOE2 or APOE3. Interestingly, the APOE4-associated perturbation in the female bone proteome was more pronounced than the corresponding alterations observed in the hippocampus. APOE4 protein causes bone fragility in females, but not males, even without changes in cortical bone structure. These bone quality deficits arose from suppression of osteocyte perilacunocanalicular remodeling. We find that APOE4 is a new molecular culprit capable of disrupting osteocyte maintenance of bone quality as early as midlife in a manner that disproportionately affects females. These findings highlight osteocytes as potential targets for early diagnosis of age-related cognitive impairment, and treatment for bone fragility, in females.

## INTRODUCTION

Age-related multimorbidity poses a formidable and growing public-health burden. Epidemiological data consistently demonstrate that individuals with bone fragility and osteoarthritis experience an increased incidence of cognitive impairment, Alzheimer’s disease (AD), and related dementias ^1-4^. Reciprocally, individuals diagnosed with AD are at 2-2.5 times increased risk of bone fracture than healthy individuals^5^. While these findings are true for men and women, clinical evidence supports a unique relationship between cognitive and skeletal health, specifically in women. Both AD and osteoporosis disproportionately affect females^6,7^. Post-menopausal decline in estrogen is a well-established driver of bone loss in aged females^8,9^ and is also implicated in female AD^10,11^ - although it is not the sole driver. Furthermore, a diagnosis of osteoporosis is the earliest predictor of AD, specifically in females^12^. Understanding the mechanisms responsible for the concurrent and dimorphic decline in skeletal and cognitive health with age could point to new diagnostic and therapeutic strategies for two widespread age-associated chronic diseases.

Several cellular, molecular, genetic, environmental, and lifestyle risk factors are associated with the progression of AD, including amyloidosis, Tau accumulation and aggregation, sex, exercise, diet, and many others^13,14^. Genetic variants of *APOE* in humans confer risk or protection from cognitive decline with age, with the hippocampus identified as a highly vulnerable brain region along with medial temporal regions such as the entorhinal cortex^15^. The *APOE ε4* allele is the most common and significant risk factor of late-onset AD. Relative to the neutral *APOE ε3* allele, the *APOE ε2* allele protects against AD. On the other hand, the frequency of developing late-onset AD (LOAD) with age increases from 10-20% in a non-carrier to 30-50% for *APOE ε4* heterozygotes, and 50-70% for non-Hispanic whites homozygous for *APOE ε4^16,17^*. The diverse effects generated by the differences in *APOE* allele status in humans also vary by sex, with females showing higher vulnerability to LOAD than males even in heterozygous *APOE ε4* carriers^11,18^.

While the effects of *APOE* allele status on the brain are well studied, much less is known on how the different *APOE* genotypes impact the musculoskeletal system. Since the blood-brain barrier is impermeable to circulating APOE, the brain-intrinsic and systemic functions of APOE are thought to be independent in their cellular targets, although linked in individuals with *APOE* variant alleles. Peripheral or circulating APOE, generated primarily by liver hepatocytes, influences lipid and cholesterol homeostasis and trafficking throughout the body^19-21^. People carrying *APOE ε2* and *APOE ε4* alleles have differing risks for cardiovascular disease due to changes in APOE affinity for atherogenic low density lipoprotein and triglyceride-rich, very low density lipoprotein^22-24^. In skeletal muscle, *APOE ε4* is associated with altered muscle pain perception or chronic muscle pain^25,26^. *APOE ε4* also exerts sex and diet-dependent effects on muscle resting energy expenditure by affecting mitochondrial processes^27,28^. While these findings establish a role for APOE in the musculoskeletal system, particularly in muscle, less is known about the role of APOE in bone.

More recent evidence shows that *APOE ε4* can reduce bone mineral density and increase fracture risk in humans^29^, and circulating APOE levels impact fracture healing in mice^30^. These observations, though, fail to explain why females bear the highest APOE4-associated risks or how *APOE* allele differences drive bone fragility. Importantly, bone fragility may arise from a lower quantity or quality of bone. Bone quantity is monitored clinically through bone mineral density (BMD) measurements, which are used to diagnose osteoporosis and predict fracture risk. Yet nearly half of all hip fractures in older females occur in individuals without osteoporosis or clinically low BMD^31^, underscoring the importance of bone quality—features that are independent of BMD. Although not yet assessed clinically, bone quality is influenced by geometry, microarchitecture, and material properties. Distinct cellular mechanisms regulate these traits: osteoclasts and osteoblasts, coordinated by osteocytes, control bone quantity^32-39^, while osteocytes directly govern bone quality through perilacunar/canalicular remodeling (PLR), which alters material properties that affect bone brittleness^40-44^. Osteocytes are highly dendritic, long-lived cells, that share many similarities with neurons, and compose nearly ninety percent of cells within the adult skeleton^40,45,46^. We hypothesize that osteocytes share with hippocampal neurons a unique vulnerability to *APOE ε4*, which compromises bone quality, as seen in people carrying *APOE ε4*. Therefore, to better understand the relationship between AD and bone fragility, we examined the effects of human *APOE* allele variants on the skeleton in male and female humanized *APOE* knock-in mice^47^ from the molecular to functional scales, including multimodal evaluation of the bone and brain.

## MATERIALS AND METHODS

*See **Supplemental Extended Methods** for full details*.

### Mice

15-month-old male and female humanized *APOE* knock-in (*APOE2*, *APOE3*, *APOE4*; Taconic Biosciences, B6.129P2-Apoetm1(APOE*2/3/4)MaeN9)^47^ carrying homozygous human ε2, ε3, or ε4 alleles were maintained at the Buck Institute for Research on Aging (IACUC protocol #2023-0018). Animals were housed at 67–74 °F, 30–70% humidity, on a 12-h light/dark cycle with ad libitum water and irradiated chow (LabDiet 5058). At 15 months, mice were euthanized by isoflurane inhalation followed by cervical dislocation. Wild-type C57BL/6 mice from the NIA aging colony (male and female, 4 and 21 months old) were maintained under specific pathogen-free conditions at UCSF (IACUC protocol #AN206686) with a 12-h light/dark cycle. Mice were anesthetized with 87.5 mg per kg ketamine and 12.5 mg per kg xylazine and transcardially perfused with 25ml ice-cold phosphate-buffered saline before decapitation and tissue collection.

### RNA Isolation, Sequencing, and Processing

Humeri from *APOE2/3/4* mice (n = 4/allele) were dissected, cleared of soft tissue and periosteum, and trimmed at the metaphysis. Marrow was removed by centrifugation (1,000 × g, 1 min). Bones were snap-frozen in liquid nitrogen, homogenized in ice-chilled QIAzol (Qiagen, 79308) using a rotor–stator homogenizer (Omni GLH), and stored at −80°C. RNA was extracted from osteocyte-enriched cortical bone with the miRNeasy Mini Kit (Qiagen, 74106) including on-column DNase digestion^48,49^. Concentration and quality were assessed by NanoDrop spectrophotometer. Libraries were prepared from 250 ng of RNA with RQN > 6 (Agilent Fragment Analyzer, DNF-472) using Universal Plus mRNA with NuQuant reagents (TECAN, 0520) through PCR amplification for 17 cycles. Libraries were pooled, quantified (Fragment Analyzer DNF-474), diluted to 1 nM, quality-checked on an Illumina MiniSeq (FC-420-1001), normalized by protein-coding read content, and sequenced on a NovaSeq X Plus (1.5B Reagent Kit, 100 cycles) to ∼30 M 50 bp paired-end reads/library^48^. Reads were aligned to the Ensembl mouse reference genome (GRCm38.78) using STAR v2.5.2b^50^.

### RNA-seq Differential Gene Expression Analysis

Differential expression was performed with DESeq2 (v1.46) in R (v4.4.3) using raw read counts^51^. DESeq2 normalized counts by estimated size factors, fit a negative binomial GLM, and calculated log₂ fold changes for pairwise comparisons (*APOE4* vs. *APOE3*, *APOE2* vs. *APOE3*, *APOE4* vs. *APOE2*). Analyses were conducted separately by sex.Sample relationships were assessed with partial least squares–discriminant analysis (mixOmics, v4.0.2; RStudio v1.3.1093)^52^, and the first two components were visualized. Differentially expressed genes (DEGs) were identified using the Wald test in DESeq2 with Benjamini–Hochberg (BH) FDR correction; genes with adjusted p < 0.1 were considered significant. DEGs from all comparisons were combined by sex, removing duplicates. Regularized log-transformed (rlog) values were z-scored for heatmap visualization (pheatmap). Hierarchical clustering was used to define co-expressed gene clusters (**Supplemental Table 1**).Functional enrichment of DEGs and gene clusters was performed with clusterProfiler (v3.21.0)^53^ using over-representation analysis of Gene Ontology biological process, molecular function, and cellular component categories, and KEGG pathways. Terms with BH-adjusted p < 0.05 were considered significantly enriched^54^ (**Supplemental Table 2**).

### Histology

Femurs from young and aged C57BL/6 mice and tibiae from *APOE2/3/4* mice (n = 4/group) were dissected, cleaned of soft tissue, fixed in 10% neutral buffered formalin for 48h, decalcified, dehydrated, and paraffin-embedded. Mid-diaphyseal axial sections were prepared for histological analyses as described previously^49,55^.

### Immunofluorescence

Immunofluorescence was performed as described^49^. Briefly, paraffin sections (n = 4/group) were rehydrated, antigen-retrieved in Unitrieve (Innovex Bio) for 1h, and blocked with Background Buster (Innovex Bio) for 30 min. Slides were incubated overnight at room temperature with primary antibodies: APOE (1:25, Abcam ab183597), cathepsin K (1:50, Abcam ab19027), sclerostin (1:50, R&D AF1589), MMP13 (1:50, Proteintech 18165-1-AP), or isotype controls (Abcam ab172730, ab37355). Secondary antibodies (Alexa Fluor Plus 594 anti-rabbit IgG, Invitrogen A32740; or anti-mouse IgG, A32742; 1:500) were applied for 2h. Sections were mounted with DAPI-containing antifade media and imaged on a Leica DMi8 confocal microscope with LAS X software. A blinded grader quantified percent antibody-positive osteocytes in cortical bone using ImageJ. Total osteocyte number was determined by DAPI-stained nuclei counts; nuclei with surrounding positive staining were considered positive. Statistical significance was assessed by two-way ANOVA with Sidak’s multiple comparison test (young vs. old) or Student’s t-test (*APOE3* vs. *APOE4*).

### Ploton Silver Nitrate Staining

The lacunocanalicular network (LCN) was visualized in tibial sections (n = 4/allele, *APOE2/3/4*) using Ploton silver nitrate staining following standard protocols as described^43,56,57^ then counterstained with Cresyl Violet. Brightfield images were acquired on a Nikon Eclipse E800 microscope. For quantification, two high-resolution fields (100×) were collected from anterior-medial, anterior-lateral, and posterior-lateral tibial regions (six images/animal). A blinded grader analyzed lacunar density and canalicular length using ImageJ. Lacunae were manually contoured and quantified with the Analyze Particles tool; canaliculi were traced from three osteocytes per image. Mean values per region were averaged across each group.

### Micro-computed Tomography (µCT)

Femurs from *APOE* mice (n = 8/allele for male, n= 6 *APOE3*, n = 9 *APOE4* female for cortical µCT and n = 4/allele, both sexes for trabecular µCT,15 months) were scanned using a Scanco µCT-50 at 10 µm voxel size. Cortical bone (50 slices) was analyzed at the tibial midshaft and trabecular bone was assessed in the metaphysis (200 µm below the growth plate, spanning 100 slices). Analyses followed the American Society for Bone and Mineral Research (ASBMR) guidelines^58^. Thresholding was performed at 30% of maximum grayscale value. Three-dimensional reconstructions were generated using Scanco software. Statistical significance was evaluated using a two-tailed Student’s t-test in GraphPad Prism (**Supplemental Tables 3-4**).

### Mechanical Testing

Left femurs (n = 9 *APOE3*, 7 *APOE4* males and n = 8 A*POE3*, 9 *APOE4* females) were hydrated in PBS and scanned by µCT prior to mechanical testing. Three-point bending was performed on an ElectroForce 3200 load frame (Bose) in the direction of primary physiological bending (posterior compression) with an 8 mm support span, preloaded to <0.2 N, and loaded at 0.01 mm/s until failure. Load-displacement data were recorded at 10 Hz. Yield load, maximum load, stiffness, post-yield displacement (PYD), and work-to-fracture were calculated from load-displacement curves using custom MATLAB scripts^37,59^. Yield was defined at a 10% decrease in stiffness; PYD was the displacement difference between yield and failure. Material properties (elastic modulus, yield stress, ultimate stress) were derived from µCT-based cross-sectional geometry (Imin/Cmin) using established equations from Turner et al. and Jepsen et al.^60,61^. Statistical analysis was performed with two-tailed Student’s t-tests (**Supplemental Table 5**).

### Proteomic Analysis – Protein Extraction

#### Cortical Bone

Femurs (n = 5/group, 21-month-old WT and *APOE2/3/4*, both sexes) were dissected, cleared of muscle and marrow, and stored at −80°C in HBSS with protease inhibitors^37,62^. Bones were briefly digested with Collagenase I/II (1:1, 1 µg/µL, 5 min, 37°C) and sonicated three times in fresh water (10min each). Demineralization was performed overnight in 1.2 M HCl at 4°C^63,64^. Demineralized bones were flash-frozen, pulverized with the Covaris CP02 cryoPREP Automated Dry Pulverizer (Covaris, Woburn, Massachusetts) and incubated in 6 M guanidine hydrochloride extraction buffer for 72h at 4°C^65^. Supernatants were collected by centrifugation (15,000 × g, 3 min) and buffer-exchanged to 10 mM Tris-HCl using 3 kDa Amicon filters.

#### Hippocampus

Frozen hippocampi (n = 5/group, female *APOE2/3/4*) were lysed in 8 M urea, 2% SDS, 200 mM TEAB, 75 mM NaCl, 1 µM trichostatin A, 3 mM nicotinamide, and HALT protease/phosphatase inhibitor. Tissues were homogenized once in a bead beater (25 Hz, 1.5 min), clarified by centrifugation (15,700 × g, 15 min, 4°C), and supernatants were collected.

### Protein Digestion

Protein concentration was determined using BCA assays. Bone (50 µg) and hippocampal (150 µg) proteins were solubilized in 4% SDS, 50 mM TEAB, reduced with 20 mM DTT, alkylated with 40 mM iodoacetamide, acidified (1.2% phosphoric acid), and processed through S-Trap micro spin columns. Proteins were digested sequentially with Lys-C (1:300, 37°C, 2 h) and trypsin (1:25, 47°C, for 1 h followed by 37°C overnight). Proteolytic peptides were eluted, vacuum-dried, desalted (Oasis 30 mg cartridges), resuspended in 0.2% FA, and spiked with indexed retention time (iRT) standards^66^.

### Mass spectrometric Acquisitions

Peptides (400 ng) from bone were analyzed on a Orbitrap Exploris 480 mass spectrometer with a Dionex UltiMate 3000 coupled liquid chromatography (LC) system; hippocampal peptides were analyzed on Orbitrap Eclipse Tribrid mass spectrometer with the same LC system. Peptides were loaded onto an Acclaim PepMap 100 C18 trap column and separated on a 75 µm × 50 cm analytical column at 300 nL/min using optimized gradients bone: 2.5–39.2% solvent B (80% ACN, 0.1% FA) over 165 min; hippocampus: non-linear 2–50% solvent B (98% ACN, 0.1% FA) over 175 min). All samples were acquired in data-independent acquisition (DIA) mode^67-69^. Full MS spectra were collected at 120,000 resolution (AGC 3e6, max IT 60 ms, 350–1,650 m/z), and MS2 spectra at 30,000 resolution (AGC 3e6, NCE 27–30, fixed first mass 200 m/z) using 26 variable DIA windows with 1 m/z overlap^69^ (**Supplementary Table 6**).

### Proteomic Data Processing and Statistical Analysis

DIA-MS data were processed in Spectronaut v17.6 using directDIA with dynamic, non-linear iRT calibration. Hippocampal data were searched against the Mus musculus UniProtKB-SwissProt proteome (86,492 entries, 01/27/2022). Bone DIA-MS data were searched against a custom spectral library (71,206 peptide entries) compiled from 150 independent bone DIA/DDA acquisitions. Trypsin/P digestion with up to two missed cleavages was specified; cysteine carbamidomethylation was fixed, methionine oxidation and N-terminal acetylation were variable. Protein identification used 1% precursor and protein q-values. Protein quantification was based on MS2 fragment ion XICs (3–6 ions), with local normalization and sparse data filtering. Differential analysis employed unpaired t-tests with Storey FDR correction^54^ (ref here). Bone proteins were considered significant if q < 0.05 with |log₂(fold-change)| > 0.58; for hippocampal proteins a threshold of |log₂(fold-change)| > 0.2 was set (**Supplemental Tables 7-10**). Partial least squares-discriminant analysis was performed in mixOmics (v4.0.2)^52^.

### Pathway and Network Analysis

Differentially regulated proteins were analyzed using QIAGEN Ingenuity Pathway Analysis (IPA). Pathway enrichment was scored by q-value, gene ratio, and z-score, indicating predicted activation or inhibition^70^. Significantly enriched pathways (corrected p < 0.05) were reported, with redundant pathways consolidated to the one with the lowest p-value. Protein co-expression networks were generated using log-normalized abundances in the Weighted Co-expression Network Analysis (WGCNA) R package^71^ as described^72^. Composite-tissue datasets were treated as a single set for module identification. Tissue-specific clusters were used for pathway enrichment via IPA as above (**Supplemental Table 11**).

### Data Availability

RNAseq data generated from mouse cortical bone enriched in osteocytes are available in the National Center for Biotechnology Information (NCBI) Sequence Read Archive (SRA) under BioProject accession number **PRJNA1314782.** Raw data and complete MS data sets were uploaded to the Mass Spectrometry Interactive Virtual Environment (MassIVE) repository, developed by the Center for Computational Mass Spectrometry at the University of California San Diego, and can be downloaded using the following link: ftp://MSV000099018@massive-ftp.ucsd.edu (MassIVE ID number: **MSV000099018**; Proteome Xchange ID: PXD068037). **[Note to reviewers]:** To access data prior during revision (before public release) please utilize the following information: Username: **MSV000099018_reviewer**, Password: **winter**.

## RESULTS

### Abundance of proteins linked to neurodegeneration in aging bone

Clinical data suggest that individuals with bone fragility and osteoarthritis are at increased risk of AD during aging while similarly, patients with AD are at risk for bone fragility and osteoarthritis^1-5^. However, the mechanisms of co-incident onset of neurodegenerative diseases and bone fragility during aging are unknown (**Figure 1A**). To investigate the age-related molecular changes in bone, we analyzed femurs from young (4-month) and aged (21-month) male and female C57BL/6 mice. Unbiased, label free mass spectrometric proteomic analysis of osteocyte-enriched (periosteum, marrow, and metaphases removed) femoral bone proteome unexpectedly revealed several proteins typically implicated in neurodegenerative disease among the top 10% most abundant proteins in bone from 21-month-old male mice from 1,928 protein groups identified (**Supplemental Table 7**). Apolipoprotein E (Apoe), apolipoprotein A1 (Apoa1), and amyloid precursor protein (App) were interspersed with proteins known to be in high abundance in bone, including collagen 1a2 and 1a2, bone sialoprotein (Ibsp), matrix extracellular phosphoglycoprotein (Mepe), and dentin matrix protein (Dmp1) among others (**Figure 1B**). Immunofluorescence on femoral cortical bone sections only detected high levels of Apoe in osteocytes of female bones from old WT mice (21-month-old), but not in young (4-month-old) bone of either sex or in old male bone (**Figure 1C**). Accordingly, we observed a significant age-dependent increase in the percentage of Apoe-positive osteocytes in females but not in males (**Figure 1D**). Importantly, at higher magnification, we detected intense staining for Apoe localized with bone-embedded osteocytes, the lacunocanalicular network, and cortical bone extracellular matrix, rather than with osteoblasts and osteoclasts (**Figure 1E**). In summary, proteins commonly linked to neurodegeneration are abundantly present in aging bone, with Apoe showing significantly higher expression in osteocytes of aged female bone compared to males.

**Figure 1:**
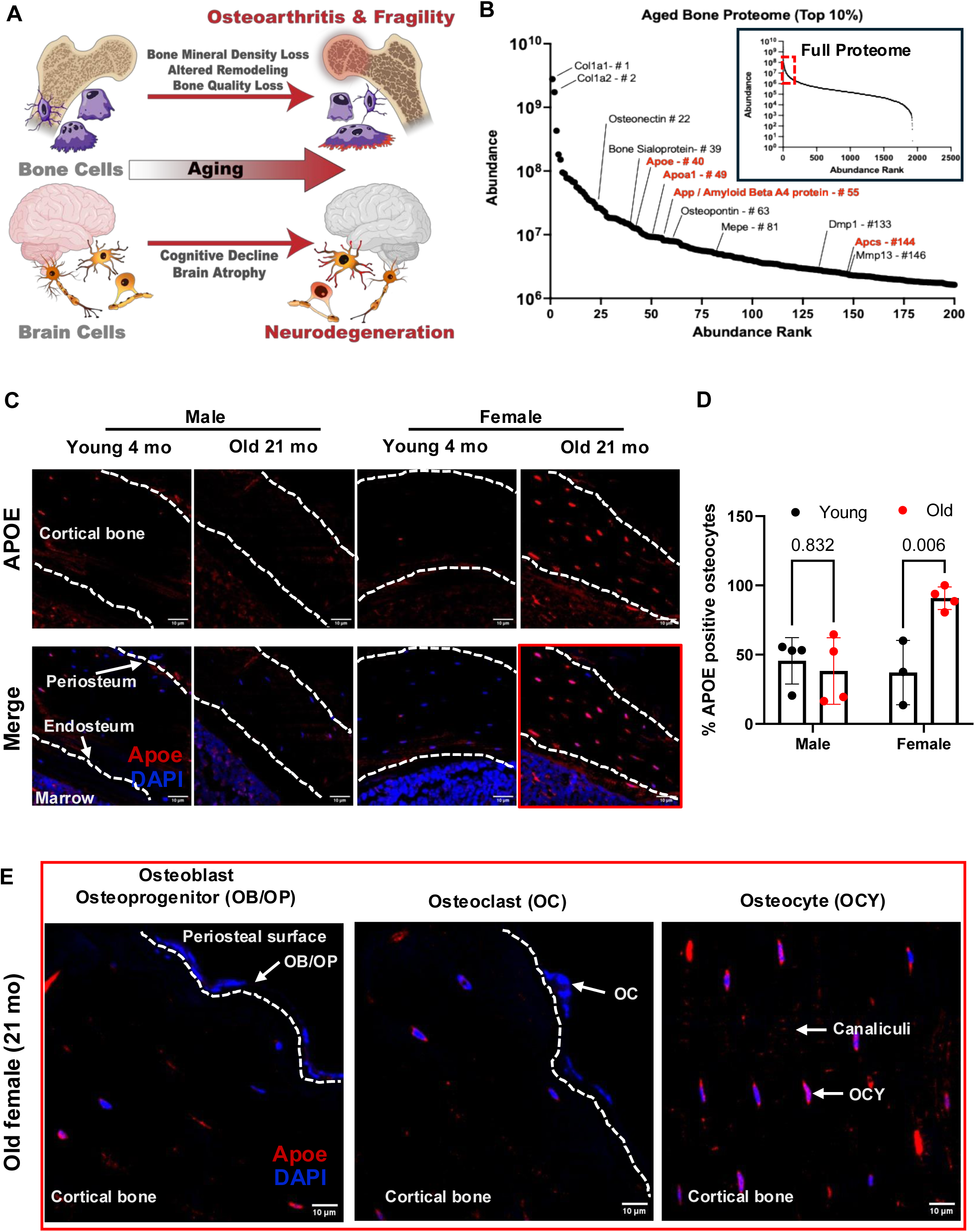
Neurological proteins are implicated in bone aging. **A**) The age-related risk for bone fragility and neurodegeneration is coupled, with tissue level deficits, such as loss of bone mineral density and brain atrophy, arising from impaired cellular function of osteoblasts, osteoclasts and osteocytes, as well as neuronal and glial cell types. With a focus on apolipoprotein E (APOE), our multimodal approach investigates the mechanisms responsible for this coordinated decline, from molecular to functional length scales. **B**) The proteome from cortical bone of aged mice (1900+ identified proteins with 2+ unique peptides per protein) shows an enrichment for common neurological and neurodegenerative proteins within the top 10% most abundant bone proteins ranked by their average abundance rank. Highly abundant proteins included neurodegenerative risk factors Apoe, amyloid precursor protein (App) the precursor to the pathogenic amyloid-β peptide, and other amyloid proteins apolipoprotein A1 (Apoa1) and amyloid P component, serum (Apcs) among common bone proteins. **C**) Immunofluorescence for Apoe expression (red) in young and old, male and female bone shows upregulation of Apoe expression with aging in female but not male bone. Scale bar: 10µm. **D**) Quantification reveals a statistically significant increase in the percentage of Apoe-positive osteocytes in aged female bone using one-way ANOVA with the p values provided above the graph. **E**) High magnification imaging demonstrating that Apoe is expressed predominantly by osteocytes in the cortical bone compared to osteoblasts and osteoclasts. Scale bar: 10µm.

### The female bone transcriptome is more susceptible to APOE allele status

Given the established role of *APOE* alleles in longevity^73-76^ and in conferring risk or protection to AD pathology, we examined whether these genetic variants similarly affect bone homeostasis. To address this question, we utilized the *APOE* knock-in mouse model harboring homozygous variants of humanized *APOE* alleles (*APOE ε2* protective, *APOE ε3* neutral, and *APOE ε4* AD risk factor) that replicate aspects of human AD in the aging mouse brain^1-3^. We performed unbiased transcriptomics and proteomics for osteocyte-enriched *APOE* bone. Transcriptomic analysis using Partial Least Square Discriminating Analysis (PLS-DA) for osteocyte-enriched bone at 15 months of age showed clear group-wise separation among the three *APOE* alleles in both male and female bone (**Figure 2A**). The female bone transcriptome was more profoundly affected by *APOE* allele status, with 146 DEGs across genotypes compared to just 22 DEGs in males (**Supplemental Table 1**). Analysis of the DEGs by hierarchical clustering revealed concordant gene expression patterns between male *APOE3* and *APOE4* bone, with opposite regulation in *APOE2* bone (**Figure 2B**). In contrast, the *APOE4* female bone drove a gene expression pattern that was distinct from that seen in *APOE2* or *APOE3* bone. These included genes involved in G-protein signaling like G protein subunit alpha O1 (*Gnao1*), which has been implicated in neurodevelopmental disorders^77^ and mitochondrial genes like ubiquinol-cytochrome-c-reductase-core-protein-2 (*Uqcrc2*). Pathway analysis identified just 2 KEGG pathways significantly enriched in male bone by *APOE* status, namely central carbon metabolism in cancer and protein digestion and absorption in the blue cluster (**Figure 2C, Supplemental Figure 1**). In female bones, KEGG pathway analysis identified 10 significantly enriched pathways, many of which are also affected by *APOE* allele status in the brain^4,5^. For example, the blue cluster shows APOE-dependent modulation of metabolic pathways including oxidative phosphorylation and thermogenesis, whereas the green cluster is enriched in the pathways dopamine synapse, serotonergic synapse, oxytocin signaling, and estrogen signaling. Interestingly, although the female red cluster contained the largest number of DE genes (85 genes; **Figure 2B**, **Supplemental Table 2**), no KEGG pathways were significantly enriched for this group. This suggests that the genes in this cluster may be functionally diverse rather than concentrated within specific KEGG pathways. However, gene ontology analysis does reveal enrichment of metabolic and neutrophil pathways in the red cluster (**Supplemental Figure 1**). In summary, *APOE* allele status is sufficient to impact the bone transcriptome, most significantly in female bone, where *APOE4* caused deregulation of metabolic, hormonal, and neurological pathways.

**Figure 2:**
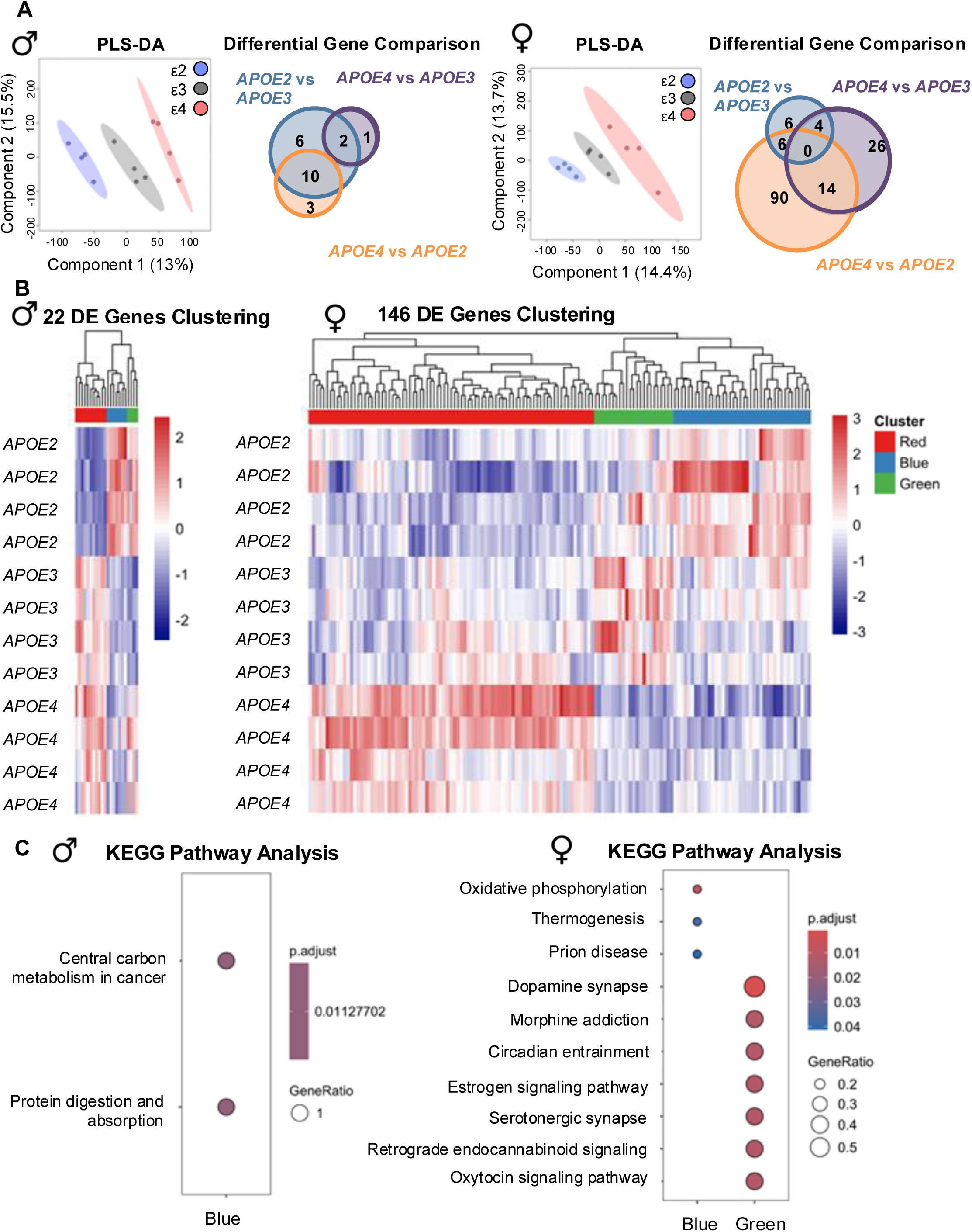
Female mouse bone transcriptome is more susceptible to the *APOE4* allele status. **A**) PLS-DA analysis of cortical bone RNAseq data exhibiting separation of groups based upon *APOE* allele status in both males and females, with Venn diagrams of DEGs in males and females showing that the female bone transcriptome is more susceptible to *APOE* allele-mediated changes than males (142 vs. 22 DEGs, FDR 0.1). **B**) Hierarchical clustering of DEGs identifies co-expressed gene clusters in males and females. **C**) KEGG pathway analysis of DEGs in each identified cluster provided no differentially regulated pathways in males except for central carbon metabolism and protein digestion and absorption in the blue cluster. However, in females, KEGG pathway analysis revealed downregulation of oxidative phosphorylation and thermogenesis in the blue cluster and downregulation of synaptic and neurobiological pathways in the green cluster.

### APOE allele status more profoundly affects the proteome of the bone than the brain

Next, we used mass spectrometry to further delineate the molecular effects of *APOE* allele status on bone. Data-Independent Acquisitions (DIA) identified 2,002 protein groups with 2 or more uniquely identified peptides per protein group from *APOE* bone samples. (**Figure 3A**). The PLS-DA showed APOE allele-dependent separation of proteomes into three groups (**Figure 3B**). Similar to the transcriptomic analysis, the differentially regulated proteins between allele pairs in female bone revealed that the largest changes were observed comparing *APOE4* mice to *APOE3* mice (393 proteins), with the fewest differentially changed proteins (19 proteins) comparing *APOE2* and *APOE3* (**Figure 3C**). All 19 of these proteins were upregulated in bones from *APOE2* mice relative to *APOE3* (**Figure 3D**). These included several complement C1q subcomponent proteins and other markers related to immune function including CD5 antigen-like (Cd5l) - also called apoptosis inhibitor of macrophage (Aim) and immunoglobulin heavy constant gamma 3 (Ighg3). The *APOE ε4* allele showed the strongest effect on the female bone proteome, whether compared to *APOE2* with 219 proteins upregulated and 32 proteins downregulated, or to *APOE3* animals with 380 upregulated and 13 downregulated proteins (**Supplemental Table 8**). In *APOE4* bones we observed consistently upregulated subunits of collagen type VI, a known marker of fibrosis^78^. Notably, the *APOE4* bone was enriched for several AD-related proteins, particularly those with known roles in amyloid-β regulation including low-density lipoprotein receptor-related protein 1 (Lrp1), Cd74, Plcg2, Lcn2, Apbb1ip, MarK2, Vcam1, Apmap, Mapk1, Ptpn1, Htra4, Csnk1a1, Cryab, and Csnk2a, which are all related to the hub gene amyloid-beta precursor protein (App) identified through IPA analysis^79^. Further, the APOE4 bone is enriched for markers of cellular senescence. In *APOE4* bones from female mice we also consistently observed downregulated markers of adaptive immunity including several immunoglobulin subunits as well as adiponectin (adipoq), which may have neuroprotective and anti-inflammatory effects in AD^80^. Proteomic analysis of male bones showed similar numbers of proteins regulated by the allele groups where *APOE4* showed distinct changes compared to *APOE2* and *APOE3*, respectively but with the directionality of regulation in males reversed when compared to females. (**Supplemental Figure 4, Supplemental Table 9**).

**Figure 3:**
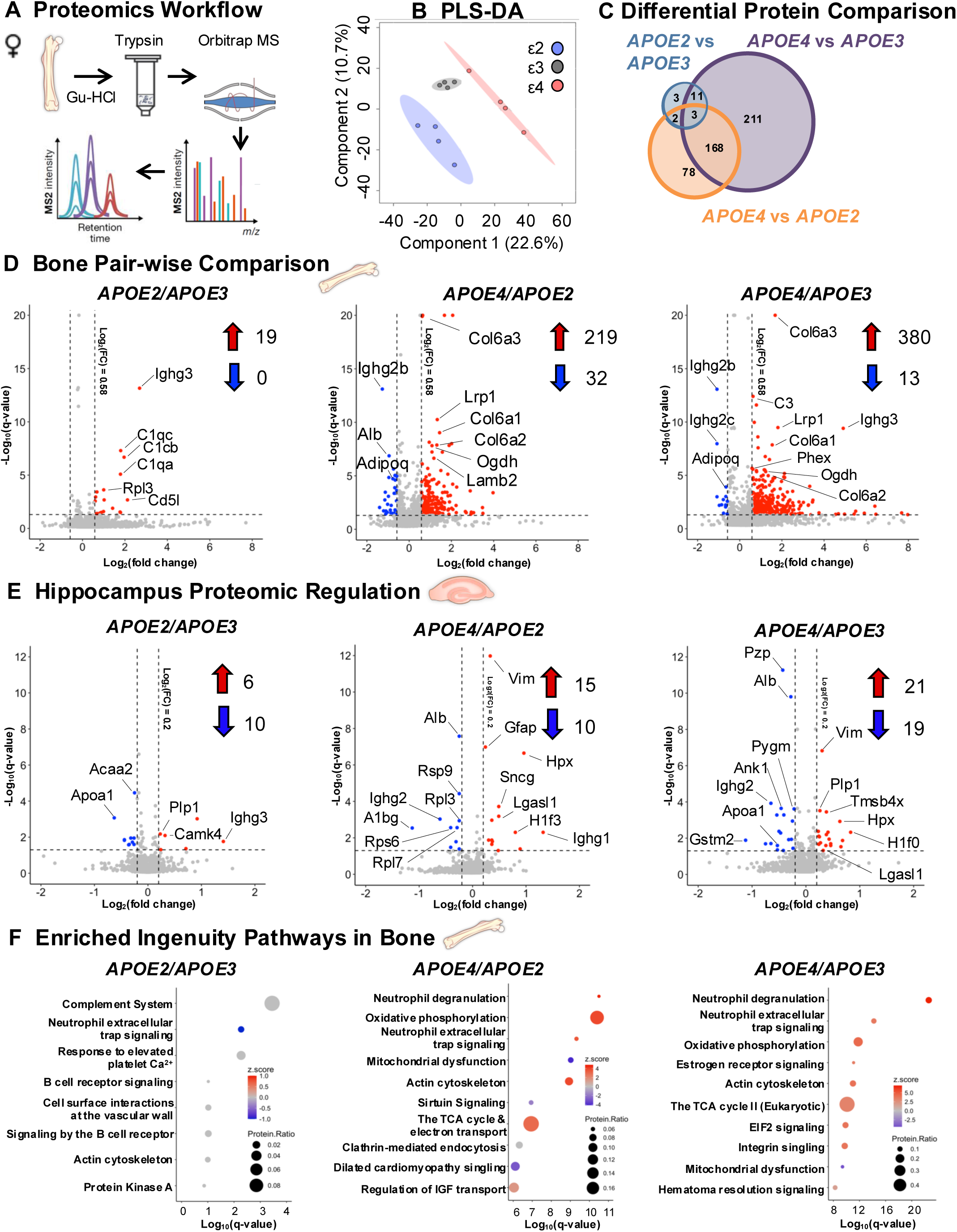
APOE4 drives differences across alleles in the bone proteome in female mice. **A**) Liquid chromatography coupled to data-independent acquisition-mass spectrometry (LC-DIA-MS) was performed on decalcified, guanidine (Gu-HCl) extracted, trypsin digested cortical bone samples from aged (15-month-old) female *APOE2/3/4* mice. Protein groups were identified (1892) with 2+ unique peptides detected per protein. **B**) PLS-DA analysis of the APOE bone proteomes showed separation of groups by *APOE* allele status. **C**) Differential pair-wise regulation of proteins by *APOE* allele (absolute Log_2_(FC) > 0.58, Q < 0.05) showed that *APOE4* drove the largest differences in protein expression when comparing genotypes. **D**) The volcano plots depict the largest number of differentially regulated proteins occurring between *APOE4* and *APOE3* where 380 proteins were significantly upregulated in APOE4 animals and only 13 proteins were downregulated. **E**) Proteomic analysis performed on the hippocampal tissue from these same animals resulted in far fewer pairwise regulated proteins, but *APOE4* consistently upregulated ECM and glycoprotein binding proteins hippocampus **F**) IPA of differentially regulated bone proteins from each pairwise comparison resulted in one predicted regulated pathway, neutrophil extracellular trap signaling, in *APOE2* vs. *APOE3*, and a mix of predicted enriched activated and repressed pathways from *APOE4* vs. *APOE2* including activation of neutrophil pathways. However, predicted pathways in *APOE4* vs*. APOE3* comparison were almost uniformly activated, including metabolic functions such as oxidative phosphorylation, electron transport, and the citric acid cycle, except the notable downregulation of mitochondrial dysfunction.

The effect of the *APOE* alleles on the bone proteome surpassed their effects on the hippocampus, a brain region heavily impacted by AD^6^. Even at a less stringent statistical threshold of Log_2_(FC) > 0.2, pairwise comparisons showed only 16 proteins regulated between *APOE2* and *APOE3*, 25 proteins regulated between *APOE4* and *APOE2*, and 40 proteins regulated between *APOE4* and *APOE3* (**Figure 3E**), and just 4, 6, and 9 proteins respectively at Log_2_(FC) > 0.58 (**Supplemental Figure 5, Supplemental Table 10**). Importantly, we had robust coverage of the hippocampal proteome, discovering over 3100 protein groups.

Surprisingly, pathways typically implicated in bone development, homeostasis, or remodeling were not among those enriched in the *APOE* bone proteome in Ingenuity Pathway Analysis (IPA). Instead, IPA identified one consistent pathway that was significantly regulated in each *APOE* allele comparison, namely, neutrophil extracellular trap signaling. While this pathway was repressed in *APOE2* relative to *APOE3* bone, neutrophil-related pathways were induced in *APOE4* relative to either *APOE2* or *APOE3* bone. In addition, the *APOE ε4* allele significantly induced or repressed several other pathways relative to *APOE2* and *APOE3* bone, including those associated with cellular metabolism (oxidative phosphorylation, TCA cycle, mitochondrial dysfunction), actin cytoskeleton and integrins, estrogen and sirtuin signaling, and others. Protein regulation in the hippocampus failed to enrich to any significantly regulated pathways in IPA analysis, but *APOE4* consistent upregulated individual ECM and glycoprotein binding proteins including vimentin (Vim), galectin-1 (Lgasl1), and hemopexin (Hpx), among others. Therefore, our results show that the APOE4-dependent female bone proteome and pathways are more broadly and comprehensively altered than in the hippocampus.

### APOE4 allele status leads to bone fragility in females due to bone quality deficits

Although *APOE* allele variants showed more profound effects on the bone proteome than on the hippocampus in 15-month-old female mice, the functional effects of these variants on bone remain unclear. To investigate the functional consequences of the APOE4-dependent changes in the bone transcriptome and proteome, we evaluated bone structural and mechanical properties using microcomputed tomography (μCT) and three-point bend testing, respectively. To focus on the alleles driving the most significant molecular changes, we compared bones from 15-month-old mice with the *APOE ε4* AD-risk allele to the *APOE ε3* neutral allele. In neither male nor female bones did μCT reveal significant differences in cortical bone structural parameters (**Figure 4A, Supplemental Table 3**). Trabecular μCT also showed no changes in male bones based on *APOE* allele status while female *APOE4* bones exhibited an increase in trabecular spacing and reduced connectivity density compared to *APOE3* controls (**Supplemental Table 4**) indicating compromised trabecular bone architecture in the absence of cortical bone structural changes.

**Figure 4:**
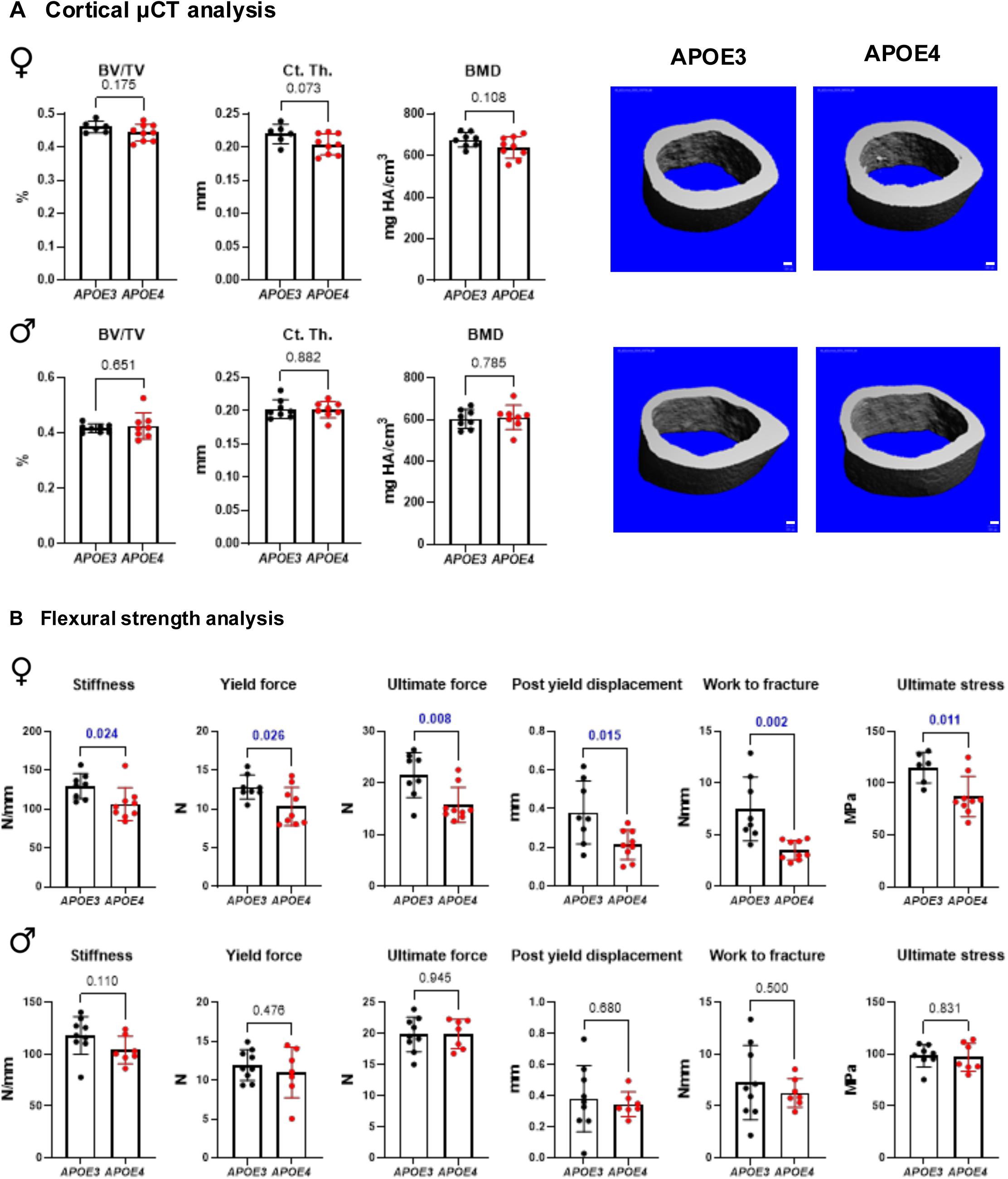
APOE4 induces bone fragility in female mice. **A**) Cortical bone μCT analysis shows no change in the bone volume fraction (BV/TV), cortical thickness (Ct. Th.), or bone mineral density (BMD) in either males or females expressing *APOE Ι4* allele compared to *APOE3* controls. Scale bar: 100µm. **B**) Flexural strength analysis of male and female femurs reveals impaired bone mechanical properties like stiffness, yield stiffness, yield force, ultimate force, post-yield displacement and work to fracture and material property of ultimate stress in *APOE4* females, while no changes in male *APOE4* bone mechanical or material properties are observed. Data are presented as mean +/- SD and statistical significance were determined with two-tailed t-test. The p-value is provided above each bar graph.

Since bone quality deficits can cause fragility, in spite of normal bone structural properties, we proceeded with three-point bend mechanical testing^7,8^. The mechanical and material properties of male *APOE3* and *APOE4* bone were indistinguishable (**Figure 4A, Supplemental Table 5**). However, in female bone, 3-point bending revealed major *APOE ε4*-dependent deficits in bone mechanical and material behavior. These deficits are apparent in multiple parameters, indicating the severe deregulation of female *APOE4* bone mechanical homeostasis. Indeed, work to fracture of female *APOE4* bones was 45% lower than *APOE3* bones, with no allele-dependent difference in male bones. The reduced ability to resist fracture results, in part, from reductions in stiffness and post yield displacement, indicating that female *APOE4* bone is both less resistant to deformation and more brittle than *APOE3 bone* (**Figure 4B**). Yield stiffness, yield force, and ultimate force are also reduced, with no changes in fracture force. In addition to mechanical properties, 3-point bending also allows measurement of material properties. Of the material properties we assessed, ultimate stress was reduced in female *APOE4* bones, accompanied by non-significant trends of decreased elastic modulus and yield stress (**Figure 4B**). These results demonstrate that the APOE4 allele is sufficient to cause bone fragility due to impaired bone quality in females but not in males.

### APOE4 induces bone fragility in females by suppressing osteocyte function

Since bone fragility in *APOE4* females is due to a bone quality deficit, and bone quality is regulated by osteocytes^7-10^, we hypothesized that osteocyte function would be suppressed in female *APOE4* bone. To test our hypothesis, we evaluated the effect of *APOE* alleles on functionally relevant osteocyte outcomes in female bone, specifically focusing on markers of perilacunar/canalicular remodeling (PLR). Histological analysis of the osteocyte lacunocanalicular network (LCN) via Ploton silver nitrate stain revealed the organized, dense LCN characteristic of intact PLR in *APOE3* bones. On the other hand, *APOE4* bones showed evidence of PLR suppression with a patchy LCN and canalicular loss (**Figure 5A-C**). Although the lacunar density was unaffected by *APOE* allele status, canalicular length was significantly reduced in *APOE4* bones compared to *APOE3*.

**Figure 5:**
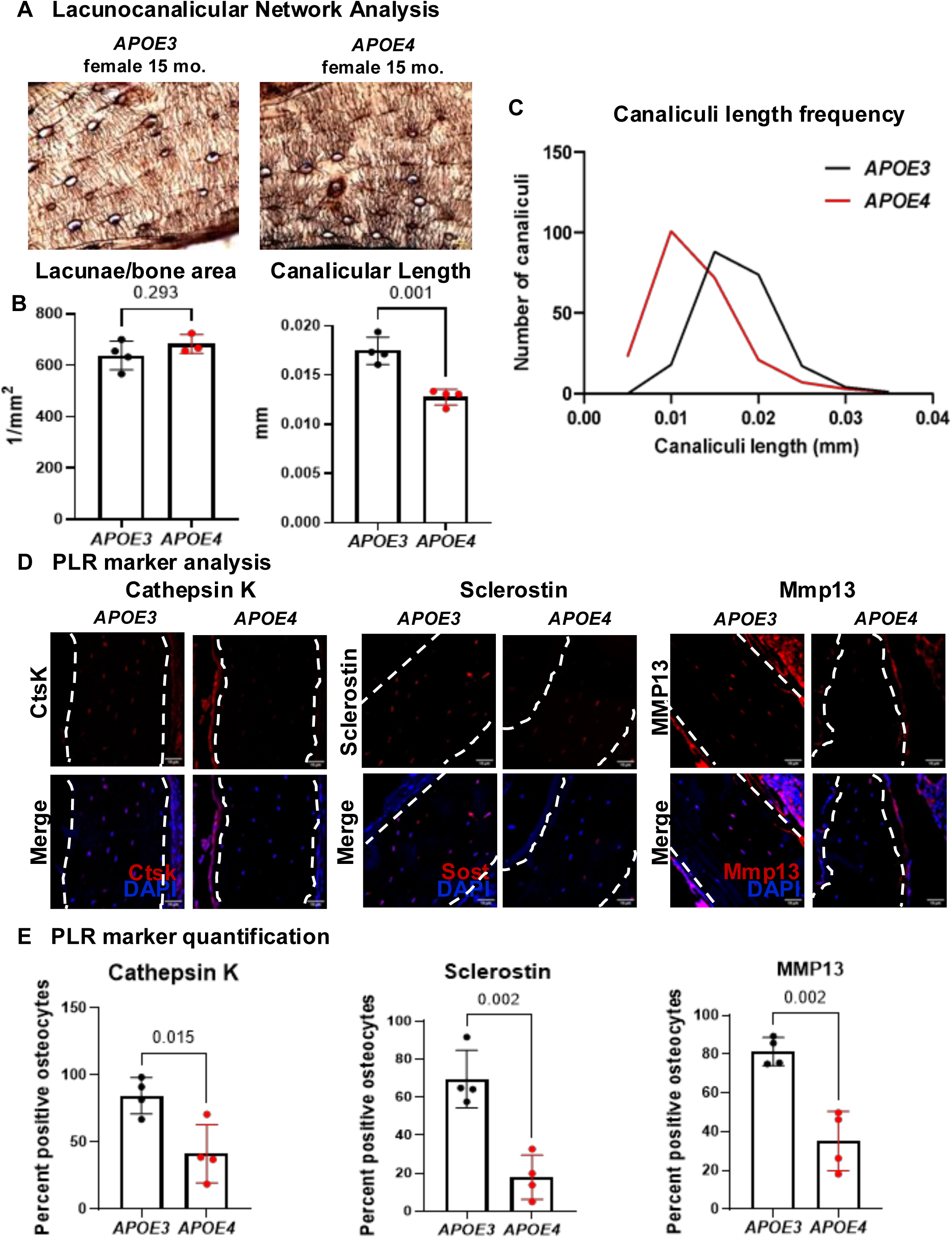
APOE4 suppresses osteocytic PLR to induce bone fragility in females. Histological analysis of female tibiae was performed to visualize the lacunocanalicular network (LCN) and analyze the expression of perilacunar/canalicular remodeling (PLR) markers. **A**) Silver nitrate staining reveals disorganized LCN and truncated canaliculi in *APOE4,* relative to *APOE3*, bones. **B**) Silver nitrate image quantification reveals a significant reduction in canaliculi length in *APOE4* bones compared to *APOE3*, with no changes in number of lacunae per bone area. **C**) The cananliculi length frequency for *APOE3* and *APOE4* bone. **D**) IF for PLR markers. and **E**) Quantification of PLR markers exhibit downregulation of cathepsin K, sclerostin and Mmp13 in *APOE4* bones compared to *APOE3*, resulting in significantly lower percent positive osteocytes in IF quantification. Scale bar: 10µm. The p-value is provided on top of each bar graph.

During PLR, mature osteocytes express cathepsin K and MMP13 to maintain LCN, a process that is accompanied by changes to osteocyte Sclerostin expression^7,11,12^. To determine if the degenerated LCN in female *APOE4* bone was accompanied by molecular evidence of PLR suppression, we analyzed PLR markers, sclerostin, cathepsin K, and Mmp13, by immunofluorescence. Although all three makers were evident in osteocytes of *APOE3* bone, sclerostin, cathepsin K, and Mmp13 were significantly reduced in osteocytes of *APOE4* bone (**Figure 5D**). Relative to *APOE3* bones, the percentage of cathepsin K, sclerostin and Mmp13 positive osteocytes in *APOE4* bones was reduced by at least 50% (**Figure 5E**). Therefore, the *APOE ε4* allele selectively causes bone fragility in females due to bone quality deficits arising from osteocytic PLR suppression.

### WGCNA Reveals Distinct but Parallel APOE Regulation in Bone and Brain

To explore whether APOEε4 regulates bone and brain through shared or distinct mechanisms, we performed Weighted Gene Co-expression Network Analysis (WGCNA) on proteomic datasets from the hippocampus and femur of 15-month-old female *APOE2/3/4* mice. Male bone was also analyzed. First, a female bone-specific WGCNA identified two protein modules significantly correlated with *APOE* allele status. These were the Brown module (155 proteins; *r* = –0.58, *p* = 0.02) and Black module (22 proteins; *r* = –0.56, *p* = 0.03) (**Supplemental Figure 4, Supplemental Table 11**). Eigengene analysis confirmed that *APOE4* mice drove group separation, with *APOE2* and *APOE3* showing similar profiles. Pathway analysis revealed expected bone-related functions (e.g., collagen synthesis, osteoarthritis) alongside many interesting neuronal and immune-related pathways, including amyloid fiber formation, cholesterol signaling, ER stress and L1cam interactions (**Figure 6B**). Enrichr analysis of the Brown module from female bone additionally predicted the mouse genomic informatics (MGI) ontology mammalian phenotype “abnormal bone remodeling” (adj p = 0.000008457), with the enriched proteins including Spon1, Sparc, Gsn, Frzb, Tnsfrsf, Ltbp3 and Gc. IPA of this cluster also predicted APP as a protein-interaction hub active in this cluster. A WGCNA for the male *APOE2/3/4* bone proteomics found no significant clusters, consistent with a lack of observed bone phenotypes (**Supplemental Figure 5, Supplemental Table 11**). Next, for WGCNA on the hippocampus we identified only one significant module (**Figure 6C, Supplemental Figure 6**, **Supplemental Table 11**), enriched in ribosomal and gene expression proteins, with minor contributions from selenium signaling, prion disease, and mitochondrial pathways. The limited brain response may reflect either distinct mechanisms or milder pathology compared to bone at this age.

**Figure 6.**
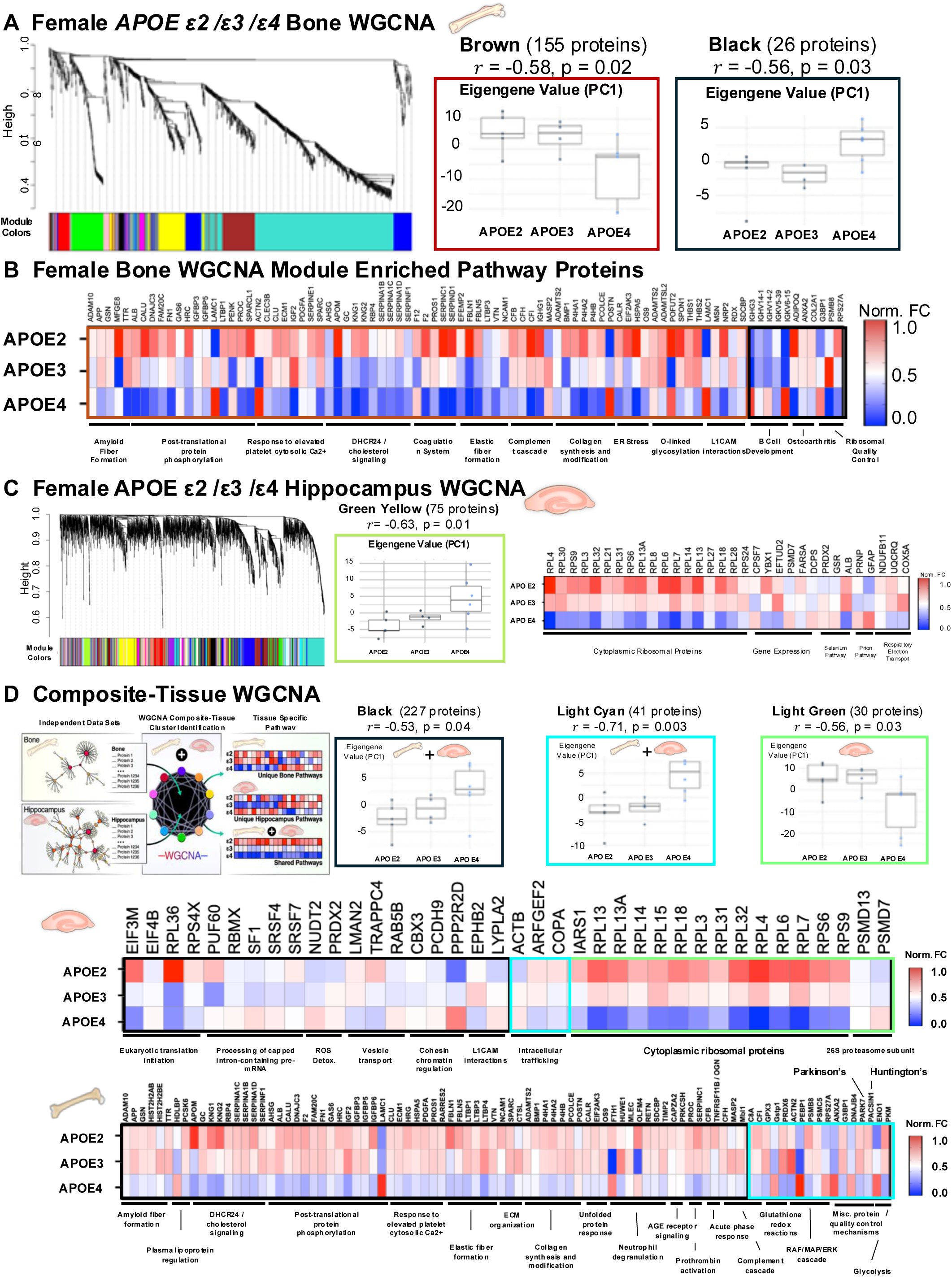
Weighted gene co-expression network analysis (WGCNA) identified APOE-dependent protein modules in female bone and hippocampus. **A**) Bone-specific WGCNA revealed two significant modules (Brown, Black). Eigengene values (principal component 1) showed allele-driven differences, primarily due to APOE4. **B**) Proteins from these clusters were analyzed by IPA, with results shown as heatmaps of allele group fold-change. Pathways included expected functions (e.g., collagen synthesis) and unexpected ones (e.g., amyloid fiber formation). **C**) Hippocampus-specific WGCNA enriched to one significant module (Green Yellow), with the strongest pathway representing cytoplasmic ribosomal proteins. **D**) Composite WGCNA combining bone and hippocampal proteomes identified modules regulated by *APOE* alleles across tissues. Significant shared modules included Black, Light Cyan, and Light Green. Black and Light Cyan clusters contained proteins from both tissues, showing APOE4-driven upregulation. The Light Green cluster, composed only of hippocampal proteins, was defined by APOE4-associated repression. IPA of hippocampal proteins from Black and Light Cyan clusters highlighted translation and mRNA processing, ROS detoxification, and vesicle transport. Parallel bone analysis revealed distinct pathways, including amyloid fiber formation, cholesterol/lipoprotein regulation, phosphorylation, ECM/collagen regulation, proteostasis, immune signaling, and wound healing. Notably, bone-specific regulation by APOE included PARK7 (Parkinson’s disease related) and PACSIN1 (Huntington’s disease related), which were not identified in hippocampal clusters in this analysis.

Finally, to assess cross-tissue regulation, we performed a composite-tissue WGCNA integrating both bone and brain proteomes. This identified 22 modules (**Supplemental Figure 7**), three of which correlated significantly with *APOE* allele: Light Cyan (41 proteins; *r* = –0.71, *p* = 0.003), Light Green (30 hippocampal proteins; *r* = –0.56, *p* = 0.03), and Black (227 proteins; *r* = –0.53, *p* = 0.04) (**Figure 6D, Supplemental Table 11**). Again, APOE4 drove group separation via eigengene analysis. This approach uncovered 79 additional APOE-regulated hippocampal proteins not seen in the hippocampus-specific WGCNA, expanding functional insights into gene translation, ROS detoxification, and intracellular transport. Notably, L1CAM interactions emerged again from the hippocampus— paralleling findings in bone—suggesting limited shared downstream regulation. In bone, the composite analysis confirmed key functions from the bone-specific WGCNA (e.g., amyloid fiber formation, ECM regulation) and revealed additional pathways, including ERK signaling, lipoprotein regulation, and neutrophil degranulation. Importantly, two neurodegeneration-associated proteins— Park7 (Parkinson’s) and Pascin1 (Huntington’s)—were newly identified as APOE-regulated in bone. Together, these findings highlight both shared and tissue-specific effects of APOE4. With the exception of L1cam signaling, APOE appears to modulate distinct downstream processes in bone and brain (**Supplemental Figures 8–9**), supporting the concept of parallel but mechanistically divergent regulation across tissues.

## DISCUSSION

The surprisingly significant and early effects of the AD risk allele *APOE4* on bone indicate that key drivers of AD may originate in the skeleton. Indeed, the scale of molecular and functional deterioration in *APOE4* female bone far exceeds that observed in the brain at the same time. Molecular dysregulation of the *APOE4* mouse proteome is more broad in bone samples than in the hippocampus. Functionally, the *APOE4* bone is nearly twice as fragile as *APOE3* controls at a time when cognitive defects are mild in these mice^81,82^. Furthermore, through mechanisms that remain unknown, Apoe accumulates with age around osteocytes in females, but not in male bone. Findings in the brain parallel the effect of *APOE4* on osteocytes, since APOE levels in the brain increase with age or disease^11,83^, and aging female brains show greater brain transcriptomic shifts than males due to APOE expression and neurodegeneration^84^. These experimental findings in mice both mirror and explain the clinical observation that osteoporosis is the earliest and strongest predictor of later AD diagnosis in women, with the osteoporosis diagnosis preceding the AD diagnosis by an average of 7 years^12^. By uncovering cellular, molecular, and mechanical mechanisms of *APOE4* action in bone, this work reveals osteocytes as untapped targets for diagnostics and interventions in both age-related cognitive and skeletal disease.

The urgent need for diagnostics and therapies to predict, prevent, or mitigate AD has driven focused research to uncover causal factors and mechanisms, largely focused on the brain. These efforts yielded powerful insight but fell short of the goal of effective clinical intervention in this devastating disease. This current study highlights the skeleton as an opportunity to unlock novel mechanisms of AD. Several factors, explored below, explain why the role of bone in AD has been invisible for so long, and advance our understanding of mechanisms of disease. First, mineralized bone presents challenges to molecular and cellular analyses. This has limited the application of the proteomic strategies that reveal the abundance of AD-related proteins in aged bone. Second, the effect of *APOE4* on bone is only apparent in female mice and women^12^, so studies that failed to examine females, or grouped both sexes, would not have detected it. Third, *APOE4* compromises osteocyte control of bone quality, while clinical and scientific efforts have largely focused on osteoblast and osteoclast control of bone quantity, which are unaffected by *APOE4* in mice. Finally, the critical endocrine and systemic role the skeleton is often overshadowed by its mechanical function. While many questions remain, available clinical and experimental evidence strongly motivate additional investigation, with the prospect that – at least in females - bone may hold the key to early diagnosis and therapeutic intervention in AD.

The surprising abundance of AD-associated transcripts and proteins in our unbiased multi-omic analyses of aged cortical bone motivated our study of AD in bone, which revealed potential mechanisms by which *APOE4* causes co-incident onset of bone fragility and neurodegenerative diseases during aging. The robust bone proteomic protocols that we developed^62^ profile bone with unparalleled depth to uncover new molecular markers in this disease model. Molecular signatures from both transcriptomic and proteomic approaches focused our attention on APOE4 specifically in female bone, such as the downregulation of metabolic functions and oxidative phosphorylation and the involvement of estrogen signaling. The observation that *APOE* allele status impacts the female bone proteome more severely than the hippocampus is intriguing and demonstrates the importance of APOE4 for skeletal as well as neurological health in females. Pathway analysis raised the question of whether the same mechanisms are sensitive to APOE allele status in both bone and the brain, especially since osteocytes and neurons share a dendritic morphology, form head-to-toe sensory networks, and lose dendrites and mitochondrial function with age^85-87^. One shared pathway found by the composite WGCNA analysis in both organ systems was neural cell adhesion molecule L1 (L1cam) interactions. In the brain, L1cam regulates neuronal development and axon outgrowth^88,89^. Though L1cam has no documented role in osteocytes, we find that *APOE4* caused a loss of osteocyte dendricity, a finding that often accompanies PLR suppression and impaired bone quality^37,38,43,44^. Given the systemic roles for APOE in regulating lipid binding and metabolism, it will be important to determine the effects of *APOE4* on osteocyte cellular metabolism and mitochondrial function in regulating PLR in future studies. Together, the proteomic and transcriptomic analyses yielded important new insights about AD and prioritize new cellular mechanisms for future studies underlying skeletal and cognitive decline.

Sex differences are increasingly recognized as critical drivers of disease risk and progression across multiple age-related conditions where women experience a disproportionate burden^90^, including heart disease^91^, stroke^92^, AD^93^, osteoporosis^6^, and osteoarthritis^94^. Findings that osteoporosis is also a strong predictor of AD, but only in females, highlight a potential sex-specific consequences of the bone–brain axis^12^. In this study, we observed that *APOE4* induces bone fragility in female mice and not in males. Both the female bone and brain transcriptomes are more severely disrupted by APOE allele status^84^, and bone quality deficits appear only in *APOE4* female mice. Though mechanisms of *APOE4* action in female bone remain to be fully elucidated, pathway analysis highlights several possible mechanisms that exhibit dimorphic control in other contexts, chief among them estrogen receptor signaling. Estrogens regulate both brain and bone metabolism. In the brain, estrogens regulate neuroplasticity and pathways implicated in cerebral energy utilization, amyloid processing and pathology^95-98^, and neurotransmitter function. In bone, estrogens exert a broad regulatory control over bone remodeling by suppressing osteoclast-mediated resorption, promoting osteoblast and osteocyte survival, and integrating bone turnover with systemic energy and inflammatory signals^99-101^. Clinically, estrogen loss increases susceptibility to osteoporosis and AD in postmenopausal women. Interestingly, the enrichment of synaptic pathways in female *APOE4* bone RNAseq parallels findings from other AD models in the brain, where females experience more pronounced synaptic connectivity loss, which coincides with greater amyloid-β burden^102^. Additionally, enrichment of neutrophil and mitochondrial pathways in the *APOE4* bone proteome is interesting since neutrophils show sex specific interferon responses and immunometabolism^103^. A recent study revealed that neutrophils exhibit sex-specific roles in AD, where in female APOE *ε4* carriers, a unique IL-17⁺ neutrophil subset infiltrates the brain and interacts pathologically with microglia, an axis that appears absent in males^104^. Another AD risk factor, TREM2 R47H variant, is linked to altered microglial activation, heightened neuroinflammation, and severe tau pathology in brains of women and female mice^105,106^. Strikingly, this same variant also drives bone fragility in females but not males with sex hormone deficiency^107^. Furthermore, changes in mitochondrial function are observed in both brain and bone with aging. Emerging evidence reveals sex-specific metabolic disruptions in AD—including female-specific deficits in brain glucose metabolism and distinct microglial metabolic profiles^108,109^. In bone, osteoclasts display female-specific dependence on mitochondrial oxidation of fatty acids and glucose in vivo^110^. Taken together, our findings reveal that factors and pathways with sexually dimorphic effects in the AD brain also operate in bone, suggesting shared mechanisms and potential common therapeutic strategies for both AD and skeletal fragility.

This susceptibility of aged female osteocytes to APOE allele status has major functional implications for skeletal health. We observed impaired whole bone mechanical properties and altered material properties even in the absence of structural changes in cortical bone, revealing that *APOE4* compromises bone quality. Osteocytes, the central regulators of bone quality through PLR^40,43,111^, showed truncated lacunocanalicular networks and suppressed function in *APOE4* females, including downregulation of PLR mediators such as cathepsin K, sclerostin, and Mmp13. To our knowledge, this is the first evidence that *APOE4* disrupts osteocyte-driven PLR to induce bone fragility. This is particularly significant given that current osteoporosis therapies primarily target bone loss but fail to address the ∼50% of fragility fractures that arise in females without measurable deficits in bone mass^31^. By uncovering a sex-specific skeletal weakness linked to a common and well-known neurologic risk factor, APOE and its allelic isoforms, we highlight a novel mechanism of bone fragility that operates independently of bone mass and extends APOE biology beyond the brain into the skeleton. These findings underscore the need for further research to unlock the translational potential of osteocytes as diagnostic and therapeutic targets for both skeletal fragility and AD in females

Beyond their primary cognitive and mechanical roles, the brain and bone also exert systemic and endocrine functions that decline with age. Neurokines such as leptin/ghrelin, and osteokines (e.g., bone gamma-carboxyglutamic acid-containing protein (BGLAP)/osteocalcin (OCN)) act across tissues to regulate metabolic homeostasis, and their dysregulation contributes to obesity and diabetes^112,113^. Importantly, these factors also interact directly within the bone–brain axis^85-87^. Several osteokines — including OCN, osteopontin (OPN), osteoprotegerin (OPG), receptor activator of nuclear factor kappa-Β ligand (RANKL), dickkopf-1 (DKK1), fibroblast growth factor 23 (FGF23), and SOST — signal to the brain, though their specific roles and links to neurodegeneration remain unclear^114-120^. Conversely, classic neurotransmitters (acetylcholine, norepinephrine) and other neuropeptides regulate bone formation and resorption through direct innervation of bone or receptor-mediated effects on osteoblasts and osteoclasts^85,121,122^. In this study, we show that the AD risk allele *APOE4* also acts peripherally on osteocytes, impairing PLR and inducing fragility in aged females. Whether these effects are unique to APOE4 or represent a broader paradigm of neurodegenerative markers influencing bone (e.g., Tau, APP) remains to be determined. Clinical links between osteoporosis/osteoarthritis and dementias and APOE-independent AD, such as those linked to Aβ and TREM2/DAP12^123-125^, and Parkinson’s disease support a wider role for bone–brain crosstalk^12,126-128^. For example, the 5xFAD mouse model, which carries APP and PSEN1 mutations but no APOE variants, shows altered bone composition (reduced mineral crystallinity, increased glycoxidation) and decreased toughness^129^. Moreover, our findings reveal that APOE4 in bone impacts proteins central to neurodegeneration, including Park7^130^ (Parkinson’s) and Pacsin1^131^ (Huntington’s). Together, these data highlight an expanded role for reciprocal signaling between bone and brain, suggesting that cross-tissue regulation contributes to the effect of APOE4 in aging and degenerative disease^73-76^.

Our findings of this *APOE4*-linked bone fragility phenotype emerge at midlife in this model where cognitive impairments are still mild^81,82^, mirroring clinical data where osteoporosis and osteoarthritis precede and predict dementia risk^12^. This shows that bone quality deficits may arise earlier than, or parallel, neurodegenerative changes, supporting growing evidence that AD pathology may extend beyond the brain to the skeleton. Targeting osteokines or neurokines could enable early diagnosis and treatment of skeletal and neurological diseases. Current therapeutic approaches primarily target bone loss but largely neglect bone quality. The identification of bone quality as a possible early biomarker of AD and modifiable by APOE4 opens new avenues for earlier diagnosis in AD and osteoporosis. This may offer new ways to influence the course of AD and its comorbidities and provide a new molecular regulator of bone quality in women. These insights underscore the need for integrated strategies that simultaneously preserve cognitive function and skeletal aging in female populations.

## ACKNOWLEDGMENTS

The authors would like to acknowledge support from the National Institutes of Health for financial support for work related to this project, including the NIA S10OD028654 (Schilling), P01AG066591 and T32AG000266 (Ellerby), and the NIDCR R01DE019284-Alzheimer’s Disease Supplement (Alliston), NIDCR R01DE01928-11A1 (Alliston), NIAMS P30AR075055, and NIA AG077770, AG067740 (Villeda). We would also like to acknowledge UCSF Bakar Aging Research Institute for postdoctoral funding support (Kaur). The authors also thank Dr. C Yee for the invaluable support provided by training coauthors in several experimental procedures.

## DISCLAIMER

This research was supported in part by the Intramural Research Program of the National Institutes of Health (NIH). The contributions of the NIH author were made as part of their official duties as NIH federal employees, are in compliance with agency policy requirements, and are considered Works of the United States Government. However, the findings and conclusions presented in this paper are those of the author(s) and do not necessarily reflect the views of the NIH or the U.S. Department of Health and Human Services.

Dr. Birgit Schilling is on the advisory board for MOBILion Systems, Chadds Ford, PA.

## AUTHOR CONTRIBUTIONS

**CAS & GK** - Conceptualization, investigation, methodology, data curation, formal analysis, validation, visualization, and writing (original draft, review & editing). **SK, JB, & CGA** - Data curation, formal analysis, software, methodology, visualization, and writing (review & editing). **KAW, HLB, MH, NML, & GB** - data curation, project administration, supervision, writing (review & editing). **SV** – funding acquisition, investigation, project administration, resources, supervision, and writing (review & editing). **LME, BS, & TA** – conceptualization, funding acquisition, investigation, methodology, project administration, resources, supervision, and writing (original draft, review & editing).

## Glossary

Human Gene (Allele) – *APOE (APOE ε4)*

Human Protein (Allele) – APOE (APOE4)

Mouse Gene* – *Apoe*

Mouse Protein* – Apoe

Humanized APOE Knock-in Mouse – *APOE2, APOE3, APOE4 (note no epsilon (ε) notation*

*\*mice are treated as not having endogenous Apoe allele differences or are considered all genetically identical for the C57black/6 (WT) line*

## Supplemental Figures

**Supplemental Figure 1: Pathway analysis for pairwise differentially expressed genes in male and female mouse bones across *APOE* allele variants.**

Pathway analysis using GO molecular factors and KEGG pathways analysis tools shows that the female transcriptome is more adversely affected by the *APOE4* risk factor relative to males, with **A**) males exhibiting changes in immunological and metabolic pathways, and **B**) females exhibiting changes in metabolic and cytoskeletal pathways, and, most interestingly, downregulation of estrogen signaling and neurobiological pathways in *APOE4* females compared to *APOE3* in KEGG pathway analysis.

**Supplemental Figure 2: Male APOE bone proteomics.**

Liquid chromatography coupled to data-independent acquisition-mass spectrometry (LC-DIA-MS) was performed on cortical bone samples from aged 15-month-old male *APOE2/3/4* mice. **A**) PLS-DA analysis on *APOE2*, *APOE3*, and *APOE4* proteomic data showed separation of groups by *APOE* allele status. **B**) Differential protein regulation showed that the *APOE4* allele drove the largest difference between the three alleles. **C**) Volcano plots depicting pairwise regulation showed moderate changes between *APOE2* and *APOE3*, but the majority of proteins were regulated by APOE4 and downregulated in males, in contrast to upregulated in *APOE4* females. **D**) IPA of differentially regulated proteins from pairwise comparisons of *APOE4* to *APOE2* and *APOE3* mice showed consistent predicted repression by Z-score with the exceptions of predicted activation of RHOGDI signaling in *APOE4* vs. *APOE2*, and predicted activation of mitochondrial dysfunction in *APOE4* vs. *APOE3*, opposite to predicted regulation in female bones.

**Supplemental Figure 3: Hippocampus protein regulation at higher fold-change cut-off.**

The number of significant protein changes in female hippocampus of *APOE2/3/4* mice were lower at statistical cutoffs of |Log_2_(FC)| > 0.58 and Q < 0.05 compared to selected |Log_2_(FC) > 0.2 and Q < 0.05 (Figure 3), such that only 4 proteins were significantly regulated between *APOE2* and *APOE3*, 6 proteins between *APOE4 vs. APOE2*, and 9 proteins between *APOE4 vs. APOE3*.

**Supplemental Figure 4: Female bone WGCNA module trait relationships.**

Module trait relationship heatmaps from WGCNA analysis of female *APOE2/3/4* cortical bone proteomes revealed two clusters (Brown and Black) with significant Pearson correlation to *APOE* allele status. *p < 0.05

**Supplemental Figure 5: Male bone WGCNA revealed no significant modules.**

Module trait relationship heatmap from WGCNA analysis of male *APOE2/3/4* bone proteomes revealed no significant modules correlated to *APOE* allele status. One cluster (Blue) trended toward a significant relationship to *APOE* alleles at p = 0.06. *p < 0.05

**Supplemental Figure 6: Female Hippocampus WGCNA module trait relationships.**

Module trait relationship heatmaps from WGCNA analysis of female *APOE2/3/4* hippocampus proteomics revealed one cluster (Green Yellow) with significant Pearson correlation to *APOE* allele status out of 58 identified clusters. Two other clusters (Dark Slate Blue & Sky Blue) trended toward a significant relationship to *APOE* alleles at *p < 0.05.

**Supplemental Figure 7: Female bone and hippocampus composite WGCNA.**

**A**) Cluster dendrogram from the WGCNA analysis of a composite data set composed of both cortical bone and hippocampal proteins from female APOE2/3/4 mice. **B**) Module trait relationship heatmap from the composite WGCNA analysis revealed three clusters with significant Pearson correlation to *APOE* allele status (Light Green, Black, Light Cyan). *p < 0.05.

**Supplemental Figure 8: Enriched pathways in hippocampus from composite-tissue WGCNA modules.**

IPA from proteins identified in composite WGCNA modules (Light Green, Black, Light Cyan). Heatmaps of the normalized fold change of individual protein abundances across the three *APOE* alleles from identified pathways are depicted for each *APOE* allele from female brain samples.

**Supplemental Figure 9: Enriched pathways in bone from composite-tissue WGCNA modules**.

IPA from proteins identified in WGCNA modules. Heatmaps of normalized fold change of individual protein abundances across the three APOE alleles from identified pathways are depicted for each *APOE* allele from female bone samples.

## Supplemental Tables

**Supplemental Table 1: Male and female DE genes across APOE genotypes**. All DEGs from male (DEG=22) and female (DEG=146) *APOE2/3/4* cortical bone, along with cluster information.

**Supplemental Table 2: Pathway analysis of male and female transcriptome clusters**. KEGG pathway analysis of DEG clusters identified from male and female *APOE2/3/4* cortical bone.

**Supplemental Table 3: *APOE ε4* allele does not alter cortical bone structural parameters in male and female mice**. Cortical bone structural parameters measured by µCT reveal no significant changes in male or female *APOE4* bones compared to *APOE3* controls.

**Supplemental Table 4: *APOE ε4* allele causes increased trabecular spacing in female bones without altering male bone structural parameters**. Trabecular bone microarchitectural parameters measured by µCT reveal no significant changes in male *APOE4* bones compared to *APOE3* controls. In females, most *APOE4* trabecular bone microarchitectural parameters are unchanged, except for increased trabecular spacing and reduced connectivity density compared to *APOE3* controls.

**Supplemental Table 5: *APOE ε4* allele induces bone fragility in female mice with no effect on male bone mechanical properties**. Flexural strength analysis via 3-point bending reveals impaired bone mechanical and material properties in *APOE4* females compared to *APOE3* controls while no change was observed in male bones with varying APOE allele status.

**Supplemental Table 6: DIA isolation schema**. Isolation scheme of the DIA acquisition method on the Orbitrap Exploris 480 mass spectrometer and the Eclipse Tribrid mass spectrometer. The DIA precursor ion isolation scheme consisted of 26 variable windows covering the 350-1,650 m/z mass range with an overlap of 1 m/z.

**Supplemental Table 7: C57/Bl6 proteomics protein abundance**. Individual and group averages for individual proteins measured by DIA-MS/MS for aged (21-month-old) C57/Bl6 (WT) mice femurs. Male, female, and pooled sex abundances for each protein are ranked from most relative abundance to least.

**Supplemental Table 8: Female bone proteomics candidates.** All quantifiable protein groups from DIA proteomics of APOE2/3/4 female mouse cortical bone (n=5 per group for each APOE genotype APOE2/3/4)) resulted in 1892 quantifiable protein groups (with ≥ 2 unique peptides). APOE2 vs. APOE3 comparison resulted in 19 proteins significantly altered with Q value < 0.05 and absolute Log2FC ≥ 0.58. APOE4 vs. APOE2 comparison resulted in 251 proteins significantly altered with Q value < 0.05 and absolute Log2FC ≥ 0.58. APOE4 vs. APOE3 comparison resulted in 393 proteins significantly altered with Q value < 0.05 and absolute Log2FC ≥ 0.58.

**Supplemental Table 9: Female hippocampal proteomics candidates.** All quantifiable protein groups from DIA proteomics of APOE2/3/4 female mouse hippocampus tissues (n=5 per group for each APOE genotype APOE2/3/4) resulted in 3138 quantifiable protein groups (with ≥ 2 unique peptides). APOE2 vs. APOE3 comparison resulted in 16 proteins significantly altered with Q value < 0.05 and absolute Log2FC ≥ 0.2. APOE4 vs. APOE3 comparison resulted in 40 proteins significantly altered with Q value < 0.05 and absolute Log2FC ≥ 0.2. APOE4 vs. APOE2 comparison resulted in 25 proteins significantly altered with Q value < 0.05 and absolute Log2FC ≥ 0.2.

**Supplemental Table 10: Male bone proteomics candidates.** All quantifiable protein groups from DIA proteomics of APOE2/3/4 male mouse cortical bone (n=5 per group for each APOE genotype APOE2/3/4) resulted in 1525 quantifiable protein groups (with ≥ 2 unique peptides). APOE2 vs. APOE3 comparison resulted in 84 proteins significantly altered with Q value < 0.05 and absolute Log2FC ≥ 0.58. APOE4 vs. APOE2 comparison resulted in 293 proteins significantly altered with Q value < 0.05 and absolute Log2FC ≥ 0.58. APOE4 vs. APOE3 comparison resulted in 352 proteins significantly altered with Q value < 0.05 and absolute Log2FC ≥ 0.58.

**Supplemental Table 11: Weighted gene co-expression network analysis cluster proteins**. Proteins and their respective identified clusters from WGCNA for the male and female bone proteomic data sets, the female hippocampus data sets, and the composite tissue WGCNA of female bone and hippocampal data sets.

