## Supplemental Figure 1 for "Alzheimer’s disease risk factor APOE4 exerts dimorphic effects on female bone"

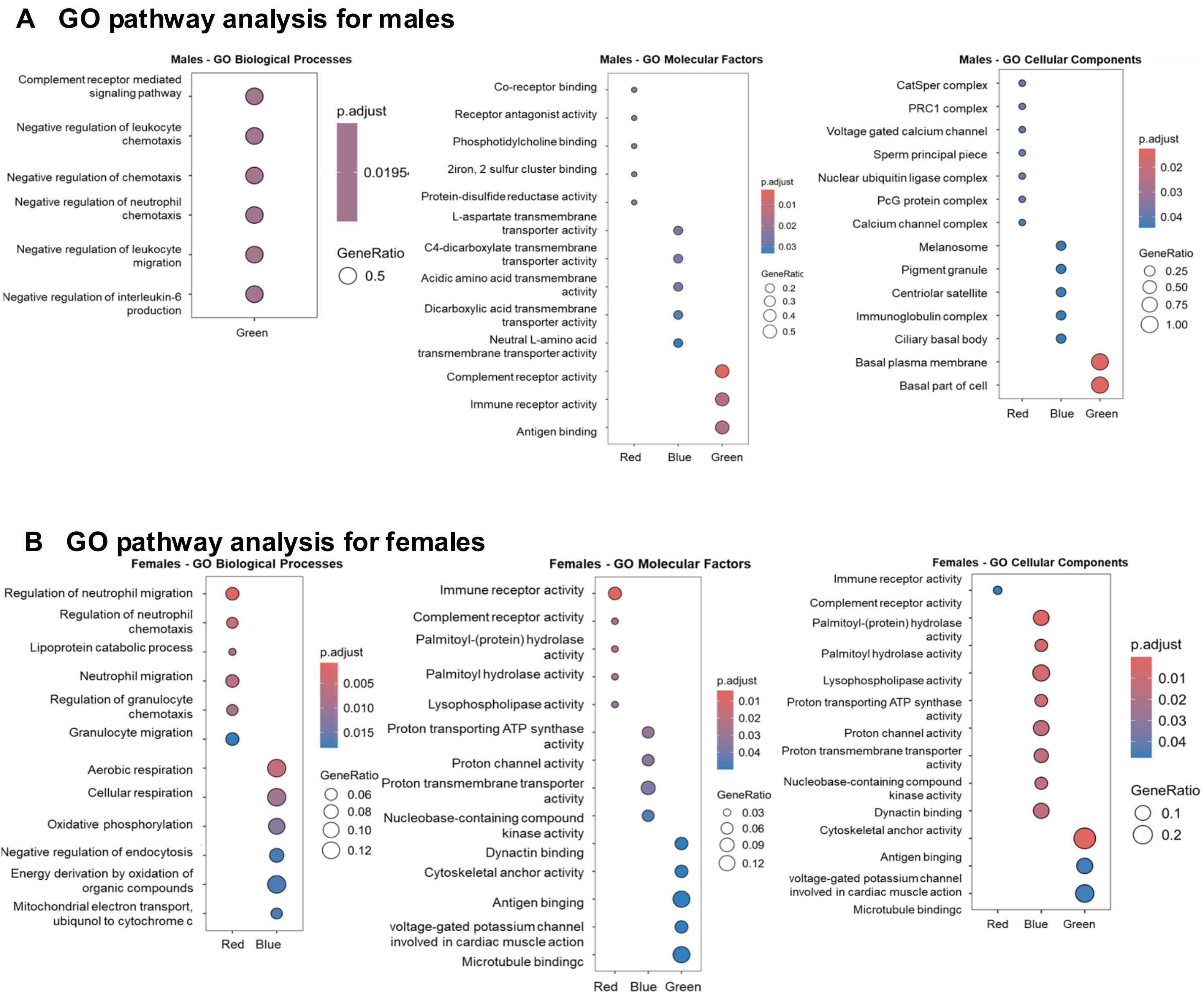


**Supplemental Figure 1: Pathway analysis for pairwise Differentially expressed genes in male and females.** Pathway analysis using GO molecular factors and KEGG pathways analysis tools shows that female transcriptome is adversely affected by the APOE4 risk factor compared to males with **(A)** males exhibiting changes in immunological and metabolic pathways while **(B)** females exhibiting changes in metabolic and cytoskeletal pathways, and, most interestingly downregulation of estrogen signaling and neurobiological pathways in APOE4 females compared to APOE3 in KEGG pathway analysis.
