## Supplemental Figure 2 for "Alzheimer’s disease risk factor APOE4 exerts dimorphic effects on female bone"

### Slide 1
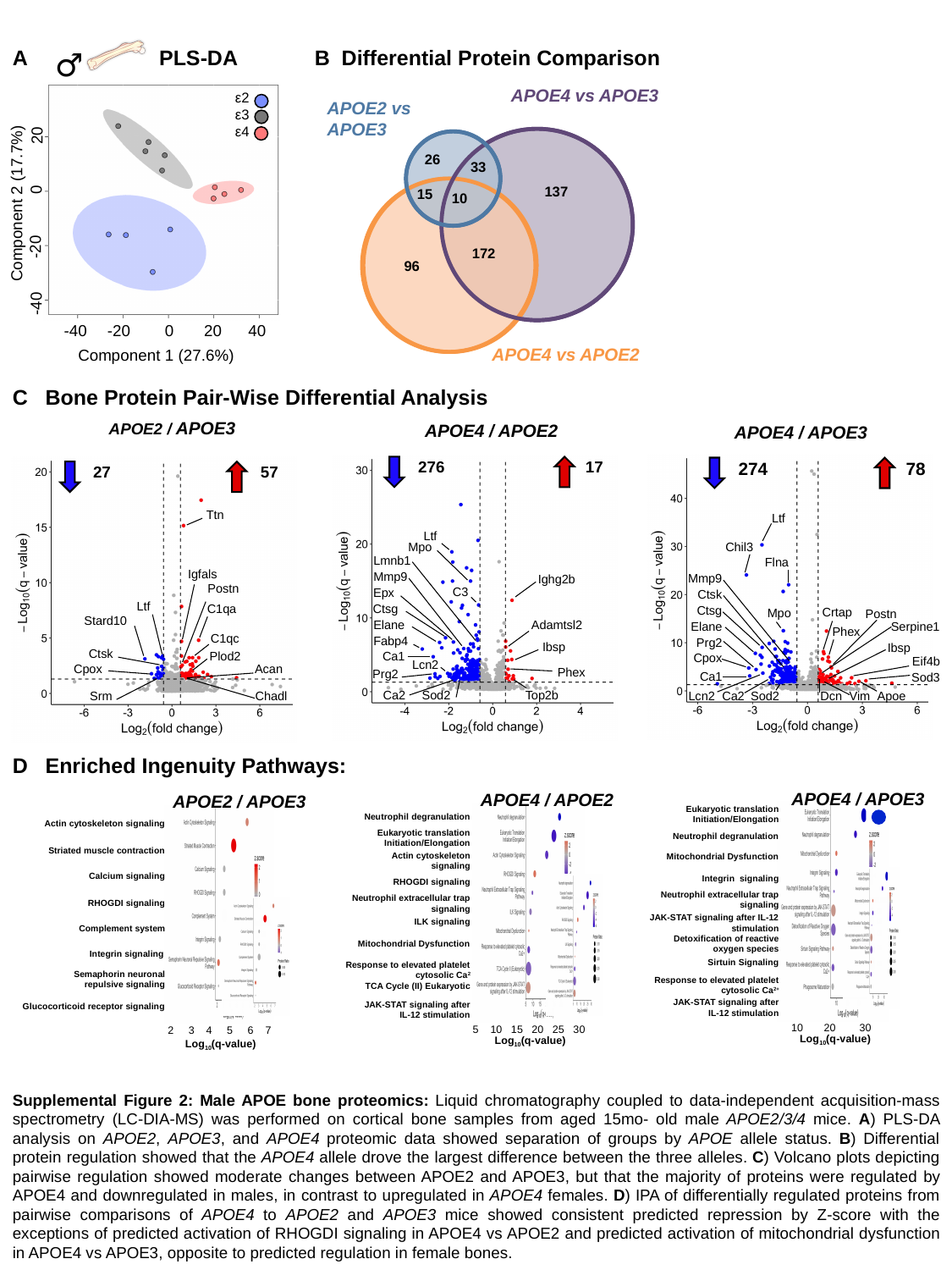

A
♂
PLS-DA
B
B Differential Protein Comparison
APOE4 vs APOE3
APOE2 vs
APOE3
26
33
137
15
10
172
96
APOE4 vs APOE2
ε2
ε3
ε4
20
0
Component 2 (17.7%)
-20
-40
-40
-20
0
20
40
Component 1 (27.6%)
C Bone Protein Pair-Wise Differential Analysis
APOE2 / APOE3
57
27
Ttn
Igfals
Postn
Ltf
C1qa
Stard10
C1qc
Ctsk
Plod2
Cpox
Acan
Srm
Chadl
APOE4 / APOE2
276
17
Ltf
Mpo
Lmnb1
Mmp9
Ighg2b
C3
Epx
Ctsg
Elane
Adamtsl2
Fabp4
Ibsp
Ca1
Lcn2
Phex
Prg2
Top2b
Ca2
Sod2
APOE4 / APOE3
274
78
Ltf
Chil3
Flna
Mmp9
Ctsk
Ctsg
Crtap
Mpo
Postn
Elane
Serpine1
Phex
Prg2
Ibsp
Cpox
Eif4b
Ca1
Sod3
Lcn2
Ca2
Sod2
Dcn
Vim
Apoe
D Enriched Ingenuity Pathways:
APOE4 / APOE3
Eukaryotic translation
Initiation/Elongation
Neutrophil degranulation
Mitochondrial Dysfunction
Integrin signaling
Neutrophil extracellular trap signaling
JAK-STAT signaling after IL-12 stimulation
Detoxification of reactive oxygen species
Sirtuin Signaling
Response to elevated platelet cytosolic Ca^2+^
JAK-STAT signaling after IL-12 stimulation
10 20 30
Log_10_(q-value)
APOE4 / APOE2
Neutrophil degranulation
Eukaryotic translation
Initiation/Elongation
Actin cytoskeleton signaling
RHOGDI signaling
Neutrophil extracellular trap signaling
ILK signaling
Mitochondrial Dysfunction
Response to elevated platelet cytosolic Ca^2^
TCA Cycle (II) Eukaryotic
JAK-STAT signaling after IL-12 stimulation
5 10 15 20 25 30
Log_10_(q-value)
APOE2 / APOE3
Actin cytoskeleton signaling
Striated muscle contraction
Calcium signaling
RHOGDI signaling
Complement system
Integrin signaling
Semaphorin neuronal repulsive signaling
Glucocorticoid receptor signaling
2 3 4 5 6 7
Log_10_(q-value)
Supplemental Figure 2: Male APOE bone proteomics: Liquid chromatography coupled to data-independent acquisition-mass spectrometry (LC-DIA-MS) was performed on cortical bone samples from aged 15mo- old male APOE2/3/4 mice. A) PLS-DA analysis on APOE2, APOE3, and APOE4 proteomic data showed separation of groups by APOE allele status. B) Differential protein regulation showed that the APOE4 allele drove the largest difference between the three alleles. C) Volcano plots depicting pairwise regulation showed moderate changes between APOE2 and APOE3, but that the majority of proteins were regulated by APOE4 and downregulated in males, in contrast to upregulated in APOE4 females. D) IPA of differentially regulated proteins from pairwise comparisons of APOE4 to APOE2 and APOE3 mice showed consistent predicted repression by Z-score with the exceptions of predicted activation of RHOGDI signaling in APOE4 vs APOE2 and predicted activation of mitochondrial dysfunction in APOE4 vs APOE3, opposite to predicted regulation in female bones.
