## Supplemental Figure 3 for "Alzheimer’s disease risk factor APOE4 exerts dimorphic effects on female bone"

### Slide 1
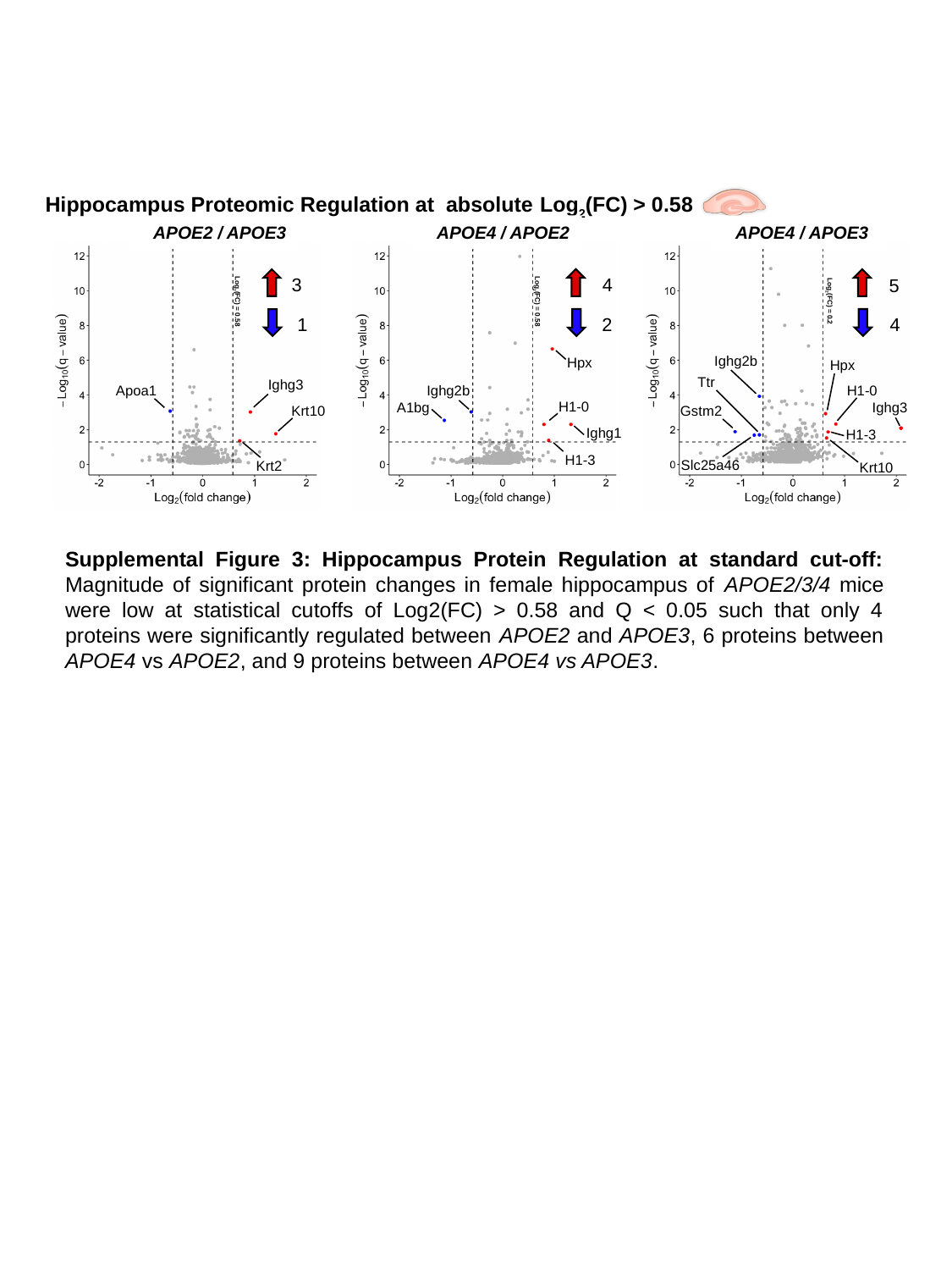

Hippocampus Proteomic Regulation at absolute Log_2_(FC) > 0.58
APOE2 / APOE3
3
1
Log_2_(FC) = 0.58
Ighg3
Apoa1
Krt10
Krt2
APOE4 / APOE2
4
2
Log_2_(FC) = 0.58
Hpx
Ighg2b
H1-0
A1bg
Ighg1
H1-3
APOE4 / APOE3
5
4
Log_2_(FC) = 0.2
Ighg2b
Hpx
Ttr
H1-0
Ighg3
Gstm2
H1-3
Slc25a46
Krt10
Supplemental Figure 3: Hippocampus Protein Regulation at standard cut-off: Magnitude of significant protein changes in female hippocampus of APOE2/3/4 mice were low at statistical cutoffs of Log2(FC) > 0.58 and Q < 0.05 such that only 4 proteins were significantly regulated between APOE2 and APOE3, 6 proteins between APOE4 vs APOE2, and 9 proteins between APOE4 vs APOE3.
