## Supplemental Figure 4 for "Alzheimer’s disease risk factor APOE4 exerts dimorphic effects on female bone"

### Slide 1
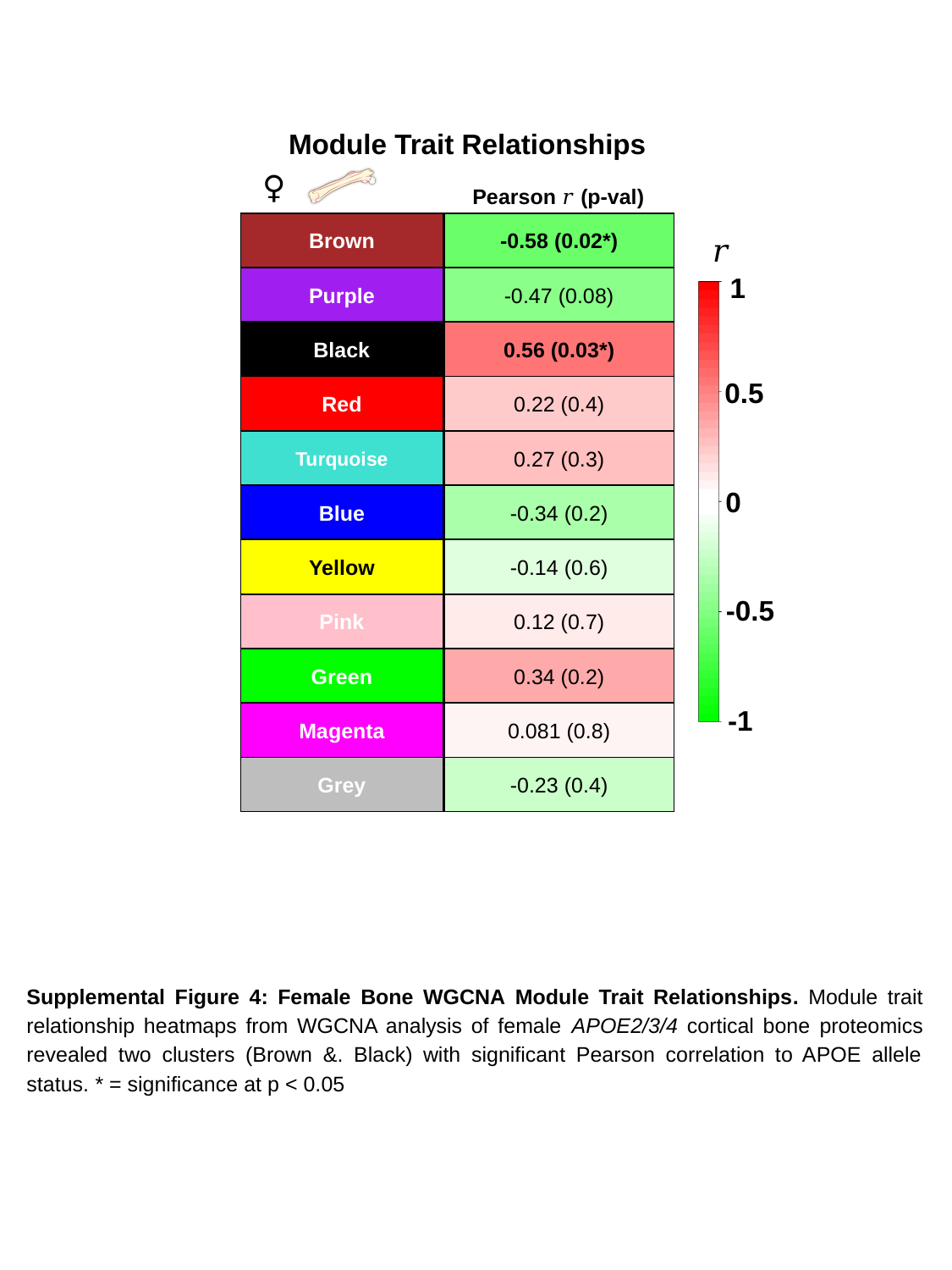

Module Trait Relationships
♀
Pearson 𝑟 (p-val)
Brown
-0.58 (0.02*)
𝑟
1
0.5
0
-0.5
-1
Purple
-0.47 (0.08)
Black
0.56 (0.03*)
Red
0.22 (0.4)
Turquoise
0.27 (0.3)
Blue
-0.34 (0.2)
Yellow
-0.14 (0.6)
Pink
0.12 (0.7)
Green
0.34 (0.2)
Magenta
0.081 (0.8)
Grey
-0.23 (0.4)
Supplemental Figure 4: Female Bone WGCNA Module Trait Relationships. Module trait relationship heatmaps from WGCNA analysis of female APOE2/3/4 cortical bone proteomics revealed two clusters (Brown &. Black) with significant Pearson correlation to APOE allele status. * = significance at p < 0.05
