## Supplemental Figure 5 for "Alzheimer’s disease risk factor APOE4 exerts dimorphic effects on female bone"

### Slide 1
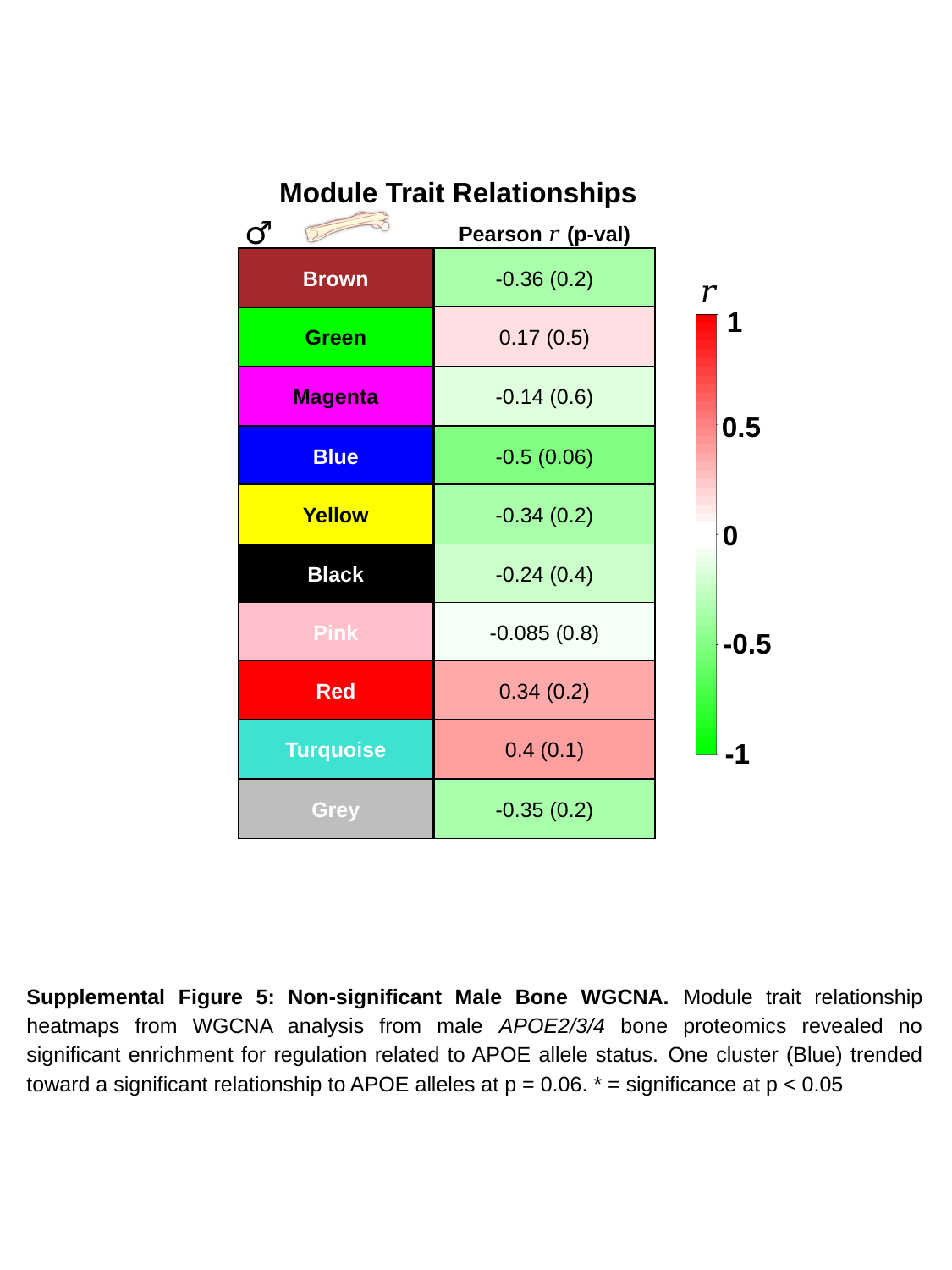

Module Trait Relationships
♂
Pearson 𝑟 (p-val)
Brown
-0.36 (0.2)
Green
0.17 (0.5)
Magenta
-0.14 (0.6)
Blue
-0.5 (0.06)
Yellow
-0.34 (0.2)
Black
-0.24 (0.4)
Pink
-0.085 (0.8)
Red
0.34 (0.2)
Turquoise
0.4 (0.1)
Grey
-0.35 (0.2)
𝑟
1
0.5
0
-0.5
-1
Supplemental Figure 5: Non-significant Male Bone WGCNA. Module trait relationship heatmaps from WGCNA analysis from male APOE2/3/4 bone proteomics revealed no significant enrichment for regulation related to APOE allele status. One cluster (Blue) trended toward a significant relationship to APOE alleles at p = 0.06. * = significance at p < 0.05
