## Supplemental Figure 6 for "Alzheimer’s disease risk factor APOE4 exerts dimorphic effects on female bone"

### Slide 1
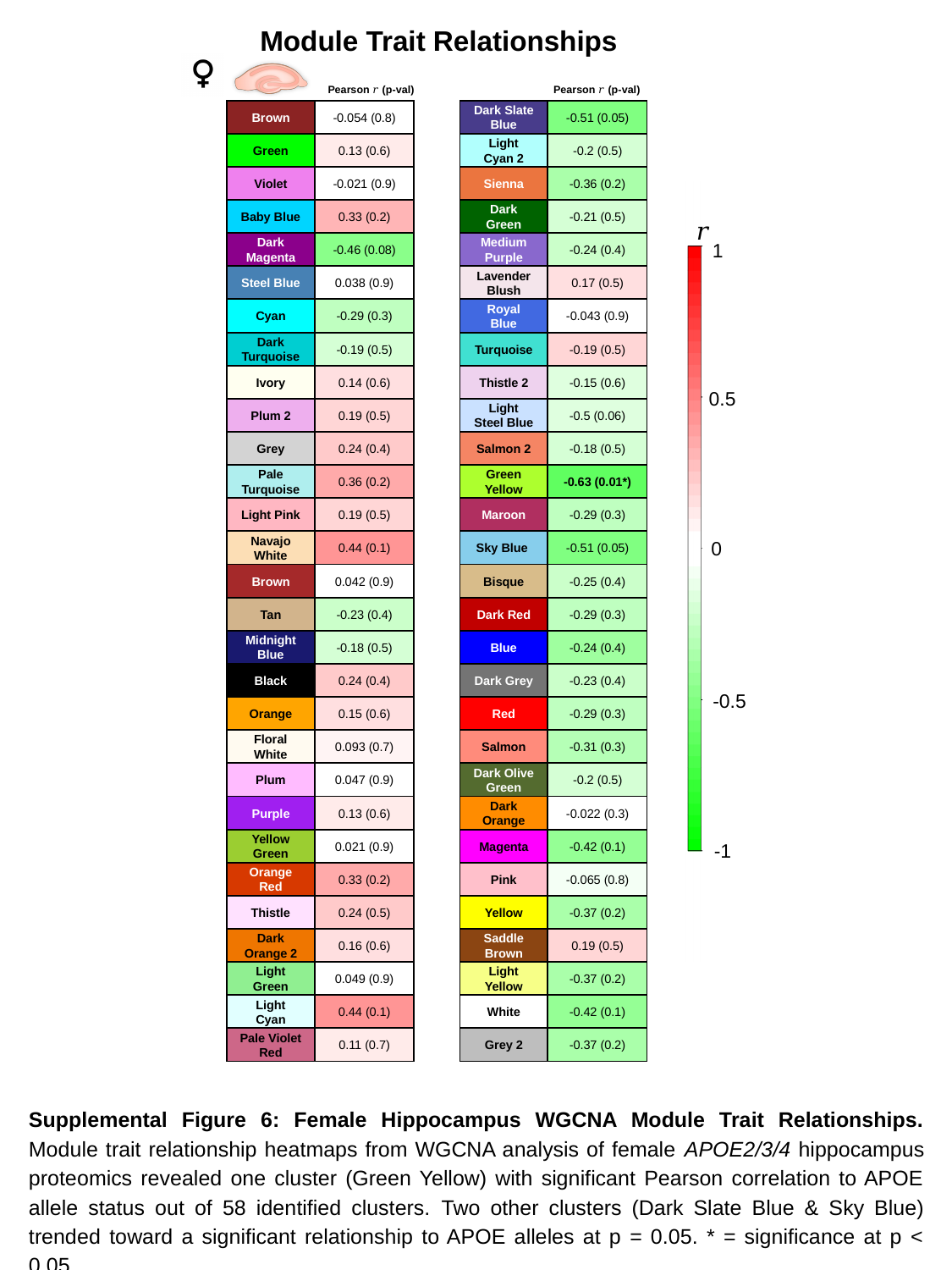

Module Trait Relationships
Pearson 𝑟 (p-val)
Pearson 𝑟 (p-val)
Brown
-0.054 (0.8)
Dark Slate Blue
-0.51 (0.05)
Green
0.13 (0.6)
Light Cyan 2
-0.2 (0.5)
Violet
-0.021 (0.9)
Sienna
-0.36 (0.2)
1
0.5
0
-0.5
-1
Baby Blue
0.33 (0.2)
Dark Green
-0.21 (0.5)
 𝑟
Dark Magenta
-0.46 (0.08)
Medium Purple
-0.24 (0.4)
Steel Blue
0.038 (0.9)
Lavender Blush
0.17 (0.5)
Cyan
-0.29 (0.3)
Royal Blue
-0.043 (0.9)
Dark Turquoise
-0.19 (0.5)
Turquoise
-0.19 (0.5)
Ivory
0.14 (0.6)
Thistle 2
-0.15 (0.6)
Plum 2
0.19 (0.5)
Light Steel Blue
-0.5 (0.06)
Grey
0.24 (0.4)
Salmon 2
-0.18 (0.5)
Pale Turquoise
0.36 (0.2)
Green Yellow
-0.63 (0.01*)
Light Pink
0.19 (0.5)
Maroon
-0.29 (0.3)
Navajo White
0.44 (0.1)
Sky Blue
-0.51 (0.05)
Brown
0.042 (0.9)
Bisque
-0.25 (0.4)
Tan
-0.23 (0.4)
Dark Red
-0.29 (0.3)
Midnight Blue
-0.18 (0.5)
Blue
-0.24 (0.4)
Black
0.24 (0.4)
Dark Grey
-0.23 (0.4)
Orange
0.15 (0.6)
Red
-0.29 (0.3)
Floral White
0.093 (0.7)
Salmon
-0.31 (0.3)
Plum
0.047 (0.9)
Dark Olive
Green
-0.2 (0.5)
Purple
0.13 (0.6)
Dark Orange
-0.022 (0.3)
Yellow Green
0.021 (0.9)
Magenta
-0.42 (0.1)
Orange Red
0.33 (0.2)
Pink
-0.065 (0.8)
Thistle
0.24 (0.5)
Yellow
-0.37 (0.2)
Dark Orange 2
0.16 (0.6)
Saddle Brown
0.19 (0.5)
Light Green
0.049 (0.9)
Light Yellow
-0.37 (0.2)
Light Cyan
0.44 (0.1)
White
-0.42 (0.1)
Pale Violet Red
0.11 (0.7)
Grey 2
-0.37 (0.2)
Supplemental Figure 6: Female Hippocampus WGCNA Module Trait Relationships. Module trait relationship heatmaps from WGCNA analysis of female APOE2/3/4 hippocampus proteomics revealed one cluster (Green Yellow) with significant Pearson correlation to APOE allele status out of 58 identified clusters. Two other clusters (Dark Slate Blue & Sky Blue) trended toward a significant relationship to APOE alleles at p = 0.05. * = significance at p < 0.05
