## Supplemental Figure 7 for "Alzheimer’s disease risk factor APOE4 exerts dimorphic effects on female bone"

### Slide 1
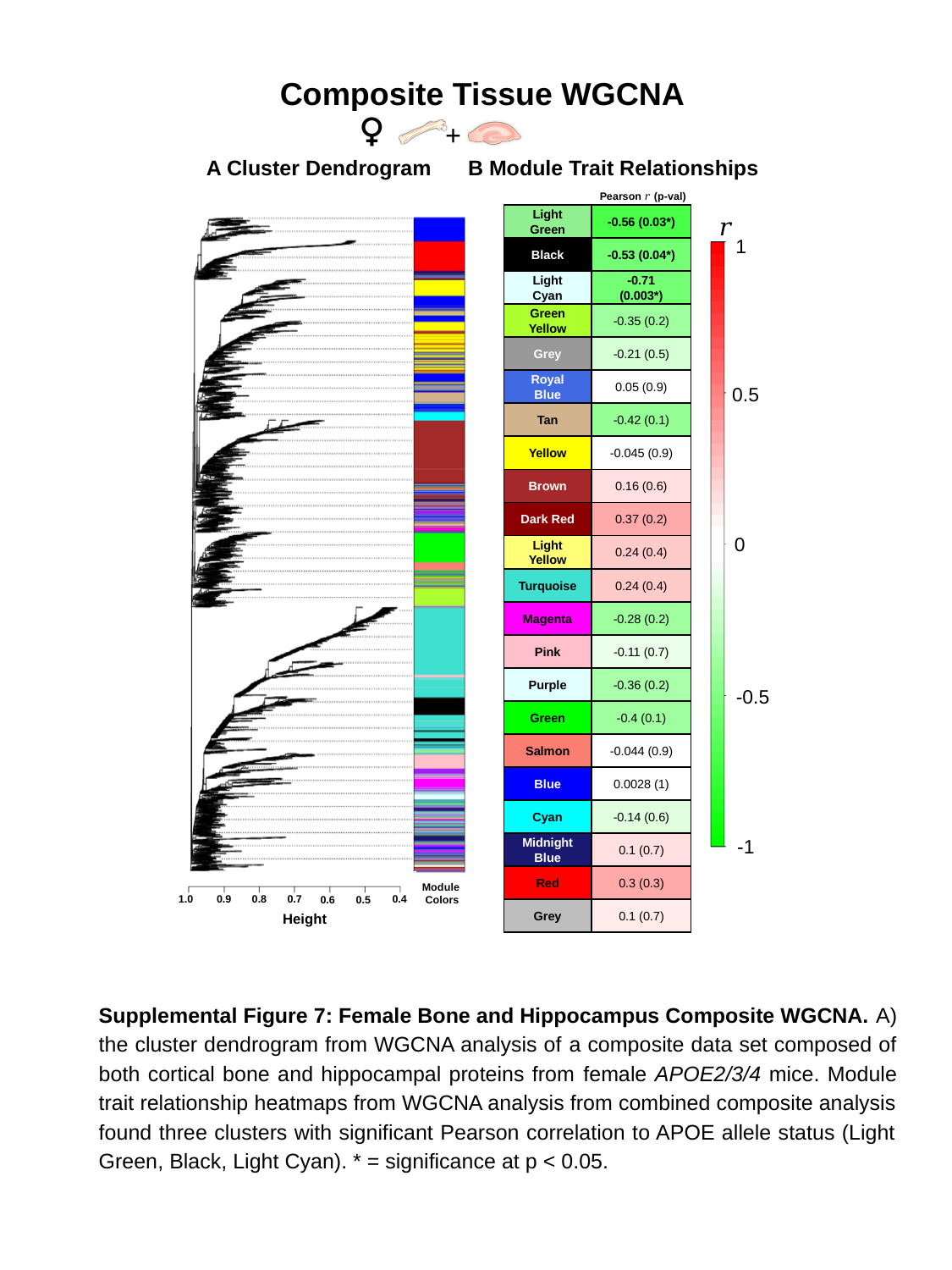

Composite Tissue WGCNA
+
A Cluster Dendrogram
B Module Trait Relationships
1
0.5
0
-0.5
-1
Pearson 𝑟 (p-val)
 𝑟
Light Green
-0.56 (0.03*)
Black
-0.53 (0.04*)
Light Cyan
-0.71 (0.003*)
Green Yellow
-0.35 (0.2)
Grey
-0.21 (0.5)
Royal Blue
0.05 (0.9)
Tan
-0.42 (0.1)
Yellow
-0.045 (0.9)
Brown
0.16 (0.6)
Dark Red
0.37 (0.2)
Light Yellow
0.24 (0.4)
Turquoise
0.24 (0.4)
Magenta
-0.28 (0.2)
Pink
-0.11 (0.7)
Purple
-0.36 (0.2)
Green
-0.4 (0.1)
Salmon
-0.044 (0.9)
Blue
0.0028 (1)
Cyan
-0.14 (0.6)
Midnight Blue
0.1 (0.7)
Red
0.3 (0.3)
Grey
0.1 (0.7)
1.0
0.9
0.8
0.7
Height
0.6
0.5
0.4
Module
Colors
Supplemental Figure 7: Female Bone and Hippocampus Composite WGCNA. A) the cluster dendrogram from WGCNA analysis of a composite data set composed of both cortical bone and hippocampal proteins from female APOE2/3/4 mice. Module trait relationship heatmaps from WGCNA analysis from combined composite analysis found three clusters with significant Pearson correlation to APOE allele status (Light Green, Black, Light Cyan). * = significance at p < 0.05.
