## Supplemental Figure 8 for "Alzheimer’s disease risk factor APOE4 exerts dimorphic effects on female bone"

### Slide 1
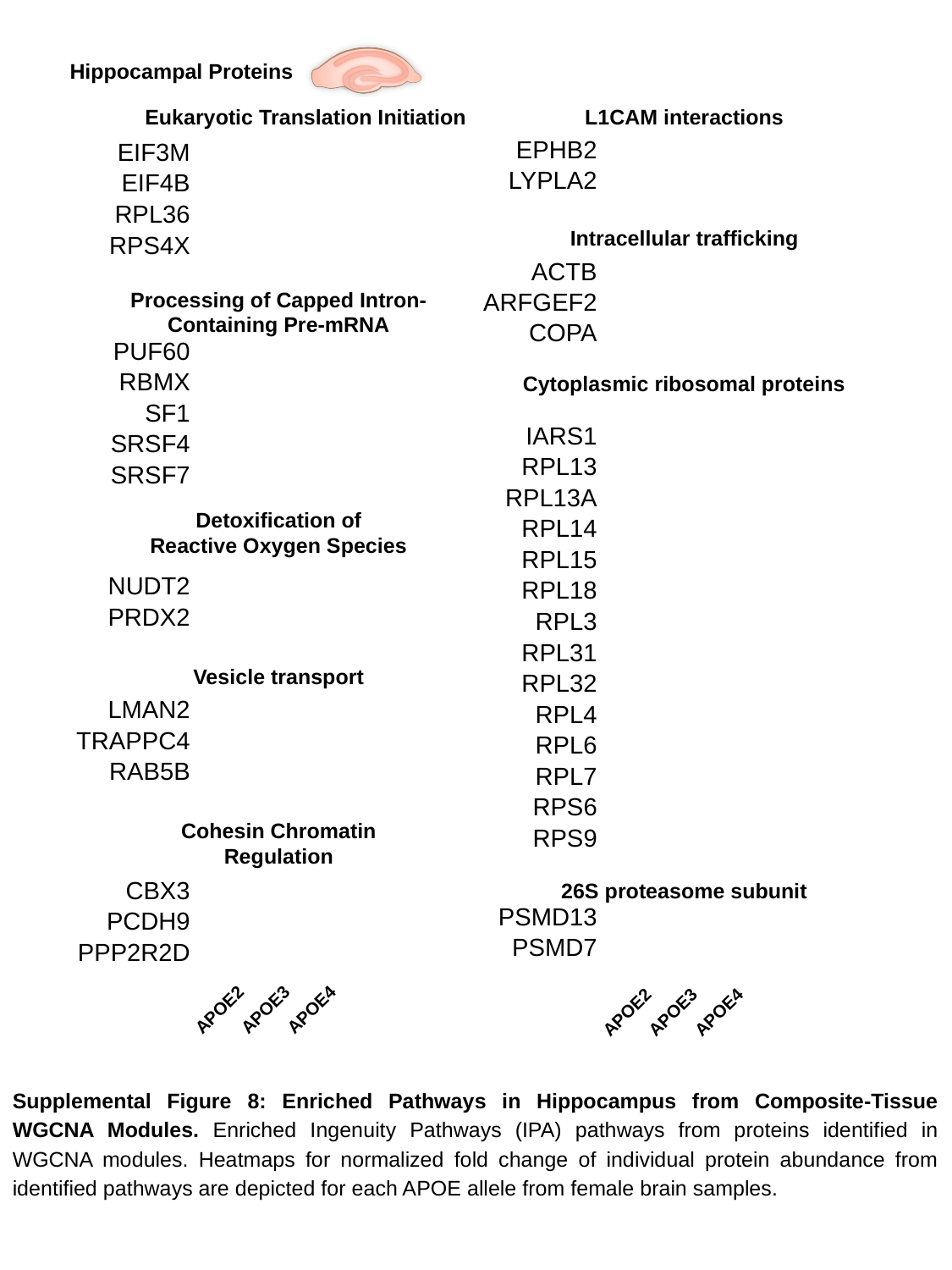

Hippocampal Proteins
Eukaryotic Translation Initiation
L1CAM interactions
| EPHB2 |
| --- |
| LYPLA2 |
| EIF3M |
| --- |
| EIF4B |
| RPL36 |
| RPS4X |
Intracellular trafficking
| ACTB |
| --- |
| ARFGEF2 |
| COPA |
Processing of Capped Intron-Containing Pre-mRNA
| PUF60 |
| --- |
| RBMX |
| SF1 |
| SRSF4 |
| SRSF7 |
Cytoplasmic ribosomal proteins
| IARS1 |
| --- |
| RPL13 |
| RPL13A |
| RPL14 |
| RPL15 |
| RPL18 |
| RPL3 |
| RPL31 |
| RPL32 |
| RPL4 |
| RPL6 |
| RPL7 |
| RPS6 |
| RPS9 |
Detoxification of Reactive Oxygen Species
| NUDT2 |
| --- |
| PRDX2 |
Vesicle transport
| LMAN2 |
| --- |
| TRAPPC4 |
| RAB5B |
Cohesin Chromatin Regulation
26S proteasome subunit
| CBX3 |
| --- |
| PCDH9 |
| PPP2R2D |
| PSMD13 |
| --- |
| PSMD7 |
APOE2
APOE3
APOE4
APOE2
APOE3
APOE4
Supplemental Figure 8: Enriched Pathways in Hippocampus from Composite-Tissue WGCNA Modules. Enriched Ingenuity Pathways (IPA) pathways from proteins identified in WGCNA modules. Heatmaps for normalized fold change of individual protein abundance from identified pathways are depicted for each APOE allele from female brain samples.
