## Supplemental Figure 9 for "Alzheimer’s disease risk factor APOE4 exerts dimorphic effects on female bone"

### Slide 1
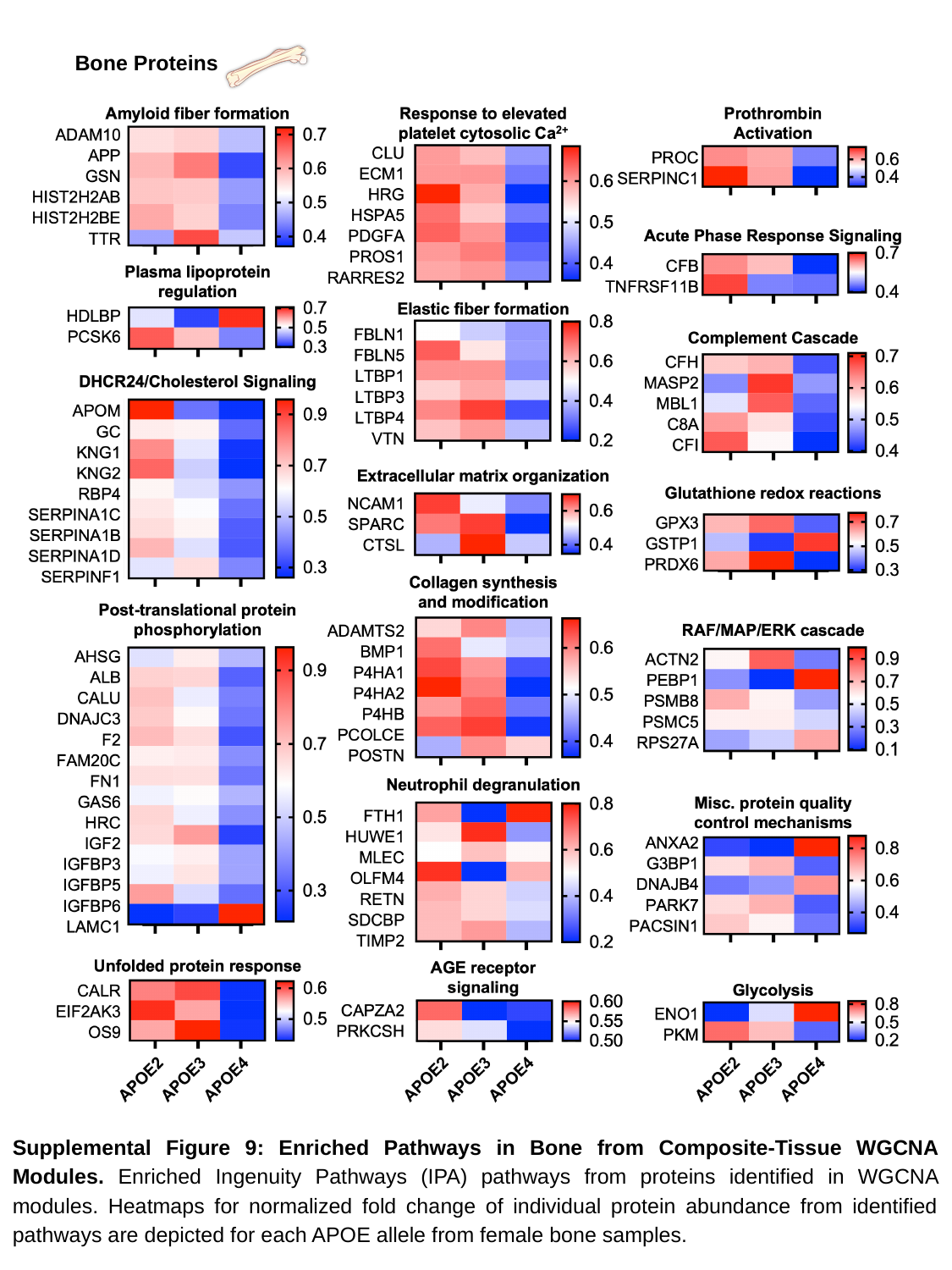

Bone Proteins
Supplemental Figure 9: Enriched Pathways in Bone from Composite-Tissue WGCNA Modules. Enriched Ingenuity Pathways (IPA) pathways from proteins identified in WGCNA modules. Heatmaps for normalized fold change of individual protein abundance from identified pathways are depicted for each APOE allele from female bone samples.
