## Supplemental Extended Methods for "Alzheimer’s disease risk factor APOE4 exerts dimorphic effects on female bone"

**EXTENDED MATERIALS AND METHODS**

**Mice**

Male and female APOE-TR mice (APOE knock-in – *APOE2/3/4*) were purchased from Taconic Biosciences B6.129P2-Apoe^tm1(APOE*2)Mae^N9 (*APOE2*), B6.129P2-Apoe^tm1(APOE*3)Mae^N9 (*APOE3*), B6.129P2-Apoe^tm1(APOE*4)Mae^N9) (*APOE4*)^1^. These animals carry homozygous copies of the human ε2, ε3, and ε4 *APOE* allele. Mice were maintained and bred in accordance with the Institutional Animal Care and Use Committee at the Buck Institute for Research on Aging using approved procedures. Mice were maintained in the mouse facility at the Buck Institute with 67°CF-74°CF, 30-70% humidity, a 12-hr light/dark cycle, and free access to water and irradiated standard chow (LabDiet 5058- PicoLab Rodent Diet 20) (IACUC Protocol # 2023-0018). 15-month-old mice were then euthanized using an IACUC-approved standard procedure of isoflurane inhalation followed by cervical dislocation.

Male and female wild type (WT) C57BL/6 mice were housed under specific pathogen-free conditions under a 12h light-dark cycle, and all animal handling and use was in accordance with institutional guidelines approved by the University of California San Francisco IACUC (Protocol # AN206686). 4-month and 21-month-old mice were then euthanized using carbon dioxide inhalation followed by cervical dislocation.

**RNA isolation and RNA sequencing and processing**

Mouse humeri (N=4 ea. *APOE2/3/4*) were dissected to remove the soft tissue and periosteum. After removing the ends of humeri at the metaphysis, the marrow was removed via centrifugation at 1000 x *g* for 1 min. Humeri were snap-frozen in liquid nitrogen, added to tubes containing ice-chilled QIAzol (Qiagen,79308), and mechanically homogenized with a rotor-stator homogenizer (GLH, Omni) followed by storage at −80 °C. mRNA was isolated from these osteocyte-enriched cortical bones using the miRNeasy Mini Kit (Qiagen, 74106) with on-column DNase digestion, following the manufacturer’s instructions as previously described^2,3^. RNA concentration and quality were quantified with a NanoDrop spectrophotometer.

RNA sequencing of bone from male and female humanized *APOE2, APOE3 and APOE4* mice was performed using NovaSeqX Plus at the UCSF Center for Advanced Technology Core^2^. RNA integrity assessed on a Fragment Analyzer (Agilent, DNF-472) with an RQN number over 6 was used for library generation. Libraries were generated from 250 ng of total RNA using Universal plus mRNA with NuQuant (TECAN, 0520) reagents through PCR amplification for 17 cycles. Individual libraries were pooled equally by volume, and quantified on a Fragment Analyzer (Agilent, DNF-474). The quantified library pool was diluted to 1 nM and further diluted as per protocol and sequenced on Illumina MiniSeq (Illumina, FC-420-1001) to check for quality of reads. Finally, individual libraries were normalized according to MiniSeq output reads, specifically by % protein coding genes, and were sequenced on NovaSeq X Series 1.5B Reagent Kit (100 cycles) (20104703) to depth of 30M reads per library. 50bp Paired-end reads were aligned to the Ensembl mouse reference genome (GRCm38.78) using STAR 2.5.2b aligner^4^.

**RNAseq differential gene expression**

Differential expression analysis was performed using the DESeq2 package (v.1.46) in R Statistical Computing Environment (v.4.4.3)^5^ with raw read counts as input. DESeq2 internally normalizes counts using estimated size factors. It fits a negative binomial generalized linear model (GLM) to the normalized counts and computes log2 fold changes in gene expression for each pairwise comparison (*APOE4* vs. *APOE3*, *APOE2* vs. *APOE3* and *APOE4* vs. *APOE2*). All analyses were performed separately for male and female mice.

To explore sample relationships and variation in gene expression, partial least squares-discriminant analysis of the transcriptomic data was performed using the package mixOmics in R (version 4.0.2; RStudio, version 1.3.1093)^6^ and the first two principal components were visualized. Differentially expressed genes (DEGs) were identified using the Wald test in DESeq2 for each pairwise comparisons. Genes with an adjusted p-value (false discovery rate, FDR) < 0.1, calculated using the Benjamini-Hochberg method, were considered significantly differentially expressed. DEGs from all three pairwise comparisons were combined separately for males and females, and duplicate genes were removed. Regularized log-transformed (rlog) values were used for heatmap visualization. Z-scores were calculated across samples for each gene, and heatmaps were generated using the pheatmap package to display gene expression patterns. Hierarchical clustering was applied to identify co-expressed gene clusters (**Supplemental Table 1**).

To investigate the biological significance of gene expression changes, functional enrichment analysis was performed using the clusterProfiler package (v.3.21.0)^7^. Over-representation analysis (ORA) was conducted on DEG lists from each pairwise comparison, as well as on gene clusters identified from heatmaps (labeled red, blue and green). Enrichment was assessed for Gene Ontology (GO) terms, including biological processes (BP), molecular functions (MF), and cellular components (CC), and Kyoto Encyclopedia of Genes and Genomes (KEGG) pathways. Terms with Benjamini-Hochberg (BH) adjusted p-values < 0.05 were considered significantly enriched^8^ (**Supplemental Table 2**).

**Histology**

Histology was conducted on femurs from young and aged C57BL/6 mice, and on tibia from *APOE* mice (n = 4 mice/group). In each case, bones were dissected to remove muscle, fixed in 10% neutral buffered formalin (NBF) for 48 hours, decalcified, dehydrated and embedded in paraffin to obtain axial sections of mid-diaphyseal bone as described before^3,9^ for various histological staining procedures described below.

**Immunofluorescence**

Immunofluorescence was performed as described^3^. Briefly, rehydrated sections (n = 4 mice/group) underwent antigen retrieval in Unitrieve (Innovex Bio, #NB325) for one hour, followed by blocking using Background Buster BB (Innovex Bio, #NB306) at room temperature for 30 minutes. The sections were then incubated overnight with either anti-APOE (1:25, Abcam, ab183597), anti-cathepsin k (1:50, Abcam, ab19027), anti-sclerostin (1:50, R&D systems, AF1589), anti-MMP13 (1:50, Proteintech, 18165-1-AP), or isotype control (rabbit: Abcam, ab172730, mouse: Abcam, ab37355) at room temperature. Subsequently, sections were incubated with secondary antibody (Alexa flour plus 594 anti-rabbit IgG (1:500, Invitrogen, A32740) or Alexa flour 594 anti-mouse IgG (1:500, Invitrogen, A32742) for two hours. The sections were then mounted using Prolong anti-fade mounting media with DAPI and visualized using a Lecia DMi8 (Leica Microsystems, Wetzlar, Germany) inverted confocal microscope operating LAS X software. ImageJ (RRID: SCR_003070) was used by a blinded grader to quantify percent positive osteocytes in the cortical bone. Cell counter feature was utilized to count number of DAPI stained nuclei in the cortical bone to obtain total osteocyte count. Next, number of nuclei surrounded with positive antibody staining was counted to calculate percent positive osteocytes number. Statistical significance was measured two-way ANOVA followed by Sidak’s multiple comparison test for young vs. old APOE IF comparisons. For APOE3/4 PLR marker quantification, two tailed Student’s t-test was utilized to evaluate significance.

**Ploton silver nitrate stain**

To visualize the lacunocanalicular network of the cortical bone, Ploton silver nitrate staining was performed as previously described^10-12^. Briefly, tibial sections (N = 4 per Apoe allele *APOE2/3/4*) were stained in a fresh mixture of 50% silver nitrate and 1% formic acid in 2% gelatin with a 2:1 ratio for 55 minutes in the dark and then counterstained with Cresyl Violet. Brightfield images of the LCN were obtained on a Nikon Eclipse E800 microscope. Two high-resolution images (100X) per tibial region (anterior-medial, anterior-lateral and posterior-lateral) were used for quantitative analysis. ImageJ (RRID: SCR_003070) was used by a blinded grader to quantify lacunar number and canaliculi length for a total of six images per animal (n=4 mice group) by converting to a binary image, manually contouring each lacunae, and measuring with the Analyze Particles feature. Canalicular length was measured by manually contouring each canaliculi of three osteocytes per image and measured. Mean values were obtained per region of the tibia per animal and were averaged within each experimental group.

**Micro-computed Tomography**

Femurs from APOE mice (n = 8/allele for male, n= 6 *APOE3*, 9 *APOE4* female for cortical µCT and n = 4/allele, both sexes for trabecular µCT,15 months) dissected from male and female mice were stored in HBSS soaked gauze at -20°C. Immediately before scanning the femurs were brought to room temperature and hydrated in 1X PBS and scanned with the following parameters: Resolution (voxel size): 10 μm, Current: 109 μA, Voltage: 55 kV, with 200 slices of epiphysis below the growth plate separation in the trabecular scan and 100 slices of mid-diaphysis in the cortical bone scan on a Scanco μCT50 (Scanco Medical, Switzerland). Cortical bone sections with automatically contoured endosteal and periosteal surfaces were analyzed using the cortical bone analysis algorithm provided in the Scanco μCT Evaluation Program. To assess trabecular parameters, including BV/TV (bone volume/total volume), Tb. N. (trabecular number), and Tb. Sp. (trabecular spacing), contours were manually segmented for 100 slices of 10 μm voxel depth starting 200μm below the growth plate and analyzed using the trabecular bone analysis algorithm provided in the Scanco μCT Evaluation Program. Analyses followed the American Society for Bone and Mineral Research (ASBMR) guidelines^13^. Statistical significance was evaluated using two tailed Student’s t-test in GraphPad Prism (**Supplemental Tables 3-4**).

**Mechanical testing**

To determine whole-bone mechanical properties, left femurs (n = 9 *APOE3*, 7 *APOE4* males and n = 8 A*POE3*, 9 *APOE4* females) subject to three-point bending in the UCSF CCMBM Skeletal Biology and Biomechanics Core immediately after μCT scanning. Load-to-failure tests were performed at room temperature using a voice coil-based mechanical load frame (ElectroForce 3200; Bose, Eden Prairie, MN) in the direction of primary physiological bending (posterior compression). Lower supports were separated by a span of 8 mm to support two ends of the specimen. The testing head was aligned at the midpoint between the supports. Femurs were preloaded to a force of <0.2 N then loaded at a rate of 0.01 mm/sec. Loading was terminated upon mechanical failure, determined by a drop in force to 0.5 N. Force displacement data was collected every 0.1 s (data rate 10 points/sec). Yield load, maximum load, stiffness, post-yield displacement (PYD), and work-to-fracture were calculated from load-displacement curves with a custom written MATLAB® script as described before^14,15^. Yield was calculated as the point where a line with a 10% decrease in stiffness intersected the force-displacement curve. PYD was calculated as the difference between the displacement at yield and the displacement at failure. Material properties of elastic modulus, yield stress, and ultimate stress were calculated from µCT measurements of left femurs using the femur cross-section diameter and moment of inertial (Imin/Cmin and Imin) and equations from Turner et al. and Jepsen et al.^16,17^ Statistical significance was evaluated using two tailed Student’s t-test in GraphPad Prism (**Supplemental Tables 5**).

**Proteomic Analysis**

**Protein Extraction**

Cortical Bone Tissue

Mouse femurs (n=5 per group for male and female for 21-month-old WT and for each APOE genotype *APOE2*/*3*/*4*) were dissected from muscle, with epiphyses removed at the growth plate and marrow removed via centrifugation to obtain diaphyseal cortical bone free of marrow and soft tissue contamination. Bones were wrapped in gauze soaked with Hank’s Balanced Salt Solution (Corning #MT21022CV) containing cOmplete, Mini, EDTA-free Protease Inhibitor (Roche #1183617000) at -80°C until preparation for Data Independent Acquisition (DIA) Mass Spectrometry (MS) analysis as described^14,18^. Briefly, to remove remaining soft tissues, thawed bones were incubated in a Collagenase I (Millipore #SRC103) and Collagenase II (Worthington #LS004177) mixture at 1:1 ratio (1 μg/μl) for 5 minutes at 37°C, followed by three rounds of sonication for 10 minutes each in fresh H_2_O. Bones were demineralized by rotating them overnight in 1 mL of 1.2 M HCl (Fischer #A144) at 4°C^19,20^. Demineralized bone was flash frozen in liquid nitrogen and mechanically pulverized using the Covaris CP02 cryoPREP Automated Dry Pulverizer (110V) (Covaris, Woburn, Massachusetts). Pulverized samples were incubated in 800 μL of extraction buffer (6 M guanidine hydrochloride Sigma #G4505, 10 mM Tris-HCl, Sigma-Aldrich #252859, 50 mM EDTA, Sigma-Aldrich #E4884) while rotating at 4°C for 72 hours^21^. Samples were spun for 3 minutes at 15,000 x g to separate pellet and soluble protein and supernatant was buffer exchanged with 500 μL of 10 mM Tris-HCl (pH 7) in Amicon 3 kDa Centrifugal Filters (MilliporeSigma #UFC900308) at 12,000 x *g* for 20 minutes three times. The final addition wash was spun until the samples were reduced to 20 μL.

Hippocampal Tissue

Frozen hippocampi from female mice (n = 5 for each APOE genotype *APOE2*/*3*/*4*) were suspended in 400 μL of lysis buffer (8 M urea, Thermo Fisher Scientific #29700, 2% SDS, Fischer #BP166, 200 mM triethylammonium bicarbonate (TEAB), Sigma-Aldrich #7408, pH 8.5, 75 mM NaCl, VWR #BDH9286, 1 μM trichostatin A, Cayman Chem #89730 , 3 mM nicotinamide, Sigma-Aldrich #N0636, and 1x HALT protease/phosphatase single-use inhibitor cocktail, Thermo Fisher Scientific #78442), and homogenized for 1 cycle with a Bead Beater Tissue Lyser II (Qiagen, Hilden, Germany) at 25 Hz for 1.5 minutes. Lysates were clarified by spinning at 15,700 x *g* for 15 min at 4°C, and the soluble protein in the supernatant was collected.

Proteolytic digestion and desalting

After protein extraction from respective tissues, protein concentration was determined using the BCA kit (Thermo Fisher Scientific #23227). 50 μg (bone) and 150 μg (hippocampal) protein samples were solubilized using 4% SDS, 50 mM TEAB. Proteins were reduced using 20 mM dithiothreitol (DTT, Sigma-Aldrich #D9779) for 10 minutes at 50°C followed by 10 minutes at RT, and alkylated using 40 mM iodoacetamide (IAA, Sigma-Aldrich #I1149) for 30 minutes at RT in the dark. Samples were acidified with a final concentration of 1.2% phosphoric acid (Sigma-Aldrich #79622) and diluted with seven volumes of S-trap buffer (90% methanol, Fischer #A452-1, in 100 mM TEAB, pH ~7). Samples were then loaded onto the S-Trap micro (bone) or mini (hippocampal) spin columns (Protifi, Farmingdale, NY, #C02-micro-80), and spun at 4,000 x *g* for 10 seconds. The S-trap columns were washed with S-trap buffer twice at 4,000 x *g* for 10 seconds each, before adding a solution of Lys-C (Roche Diagnostics, Mannheim, Germany, #11420429001) in 50 mM TEAB at a 1:300 (w/w) enzyme:protein ratio at 37°C for 2 hours. A solution of sequencing grade trypsin (Promega, San Luis Obispo, CA, #V5111) in 50 mM TEAB at a 1:25 (w/w) enzyme:protein ratio was then added, and after a 1-hour incubation at 47°C, trypsin solution was added again at the same ratio, and proteins were digested overnight at 37°C. Peptides were sequentially eluted with 50 mM TEAB (spun through for 1 min at 1,000 x *g*), 0.5% formic acid (FA, Honeywell #F0507) in water (spun through for 1 min at 1,000 x *g*), and 50% acetonitrile (ACN, Honeywell #34851) in 0.5% FA (spun through for 1 min at 4,000 x *g*). After vacuum drying, samples were resuspended in 0.2% FA in water, desalted with Oasis 30-mg Sorbent Cartridges (Waters, Milford, MA, #WAT094225). They were vacuum dried and resuspended in 0.2% FA in water at a final concentration of 1 μg/μL. Finally, indexed retention time standard peptides (iRT; Biognosys, Schlieren, Switzerland, #1900615) were spiked in the samples according to manufacturer’s instructions^22^.

**Mass spectrometric analysis**

For the protein lysates from cortical bone samples, LC-MS/MS analyses were performed on a Dionex UltiMate 3000 system online coupled to an Orbitrap Exploris 480 (Thermo Fisher Scientific, San Jose, CA). The solvent system consisted of 2% ACN, 0.1% FA in water (solvent A) and 80% ACN, 0.1% FA in water (solvent B). Digested peptides (400 ng) were loaded onto an Acclaim PepMap 100 C_18_ trap column (0.1 x 20 mm, 5 µm particle size; Thermo Fisher Scientific, # 164535) over 5 minutes at 5 µL/minute with 100% solvent A. Peptides were eluted on an Acclaim PepMap 100 C_18_ analytical column (75 µm x 50 cm, 3 µm particle size; Thermo Fisher Scientific, # 164570) at 300 nL/minute using the following gradient of solvent B: linear from 2.5% to 24.5% in 125 minutes, linear from 24.5% to 39.2% in 40 minutes, up to 98% in 1 minute, and back to 2.5% in 1 minute. The column was re-equilibrated for 30 minutes with 2.5% of solvent B, and the total gradient length was 120 minutes. For the protein lysates from hippocampal samples, LC-MS/MS analyses were performed on a Dionex UltiMate 3000 system coupled to an Orbitrap Eclipse Tribrid mass spectrometer (Thermo Fisher Scientific, San Jose, CA) with the same LC column setup and a solvent system consisting of 2% ACN, 0.1% FA in water (solvent A) and 98% ACN, 0.1% FA in water (solvent B). Peptides (400 ng) were loaded onto an Acclaim PepMap 100 C_18_ trap column over 10 minutes at 2 µL/minute with 100% solvent A. Peptides were eluted on an Acclaim PepMap 100 C_18_ analytical column at 0.3 µL/minute using the following gradient of solvent B: 2% for 10 minutes, linear from 2% to 25% in 96 minutes, linear from 25% to 40% in 23 minutes, linear from 40% to 50% in 6 minutes, up to 80% in 1 minute up, 80% for 9 minutes, and down to 2% in 1 minute. The column was equilibrated with 2% of solvent B for 29 minutes, with a total gradient length of 175 minutes. All samples, bone or hippocampus tissue, were acquired in data-independent acquisition (DIA) mode^23-25^ with consistent acquisition schemes on each instrument. Full MS spectra were collected at 120,000 resolution (AGC target: 3e6 ions, maximum injection time: 60 ms, 350-1,650 m/z), and MS2 spectra at 30,000 resolution (AGC target: 3e6 ions, maximum injection time: Auto, NCE: 27 or 30 (Eclipse or Exploris, resp.), fixed first mass 200 m/z). The DIA precursor ion isolation scheme consisted of 26 variable windows covering the 350-1,650 m/z mass range with an overlap of 1 m/z (**Supplementary Table 6**)^25^.

**DIA data processing and statistical analysis**

DIA data was processed in Spectronaut (version 17.6.230428.55965). DirectDIA searches for hippocampal data were searched against the *Mus musculus* proteome with 86,492 entries (UniProtKB-SwissProt), accessed on 01/27/2022. Cortical bone DIA analysis was searched against a custom spectral library assembled from over 150 independent acquisitions of mouse bone samples acquired in both DIA and Data Dependent Acquisition (DDA) mode composed of 71,206 peptidic entries (see *Data Availability)*. Trypsin/P was set as the digestion enzyme and two missed cleavages were allowed. Cysteine carbamidomethylation was set as a fixed modification while methionine oxidation and protein N-terminus acetylation were set as dynamic modifications. Data extraction parameters were set as dynamic and non-linear iRT calibration with precision iRT selected. Identification was performed using 1% precursor and protein q-value. For the protein level, quantification was based on the peak areas of extracted ion chromatograms (XICs) of 3 – 6 MS2 fragment ions, with local normalization and q-value sparse data filtering applied. iRT profiling was selected. Differential analysis was performed using an unpaired t-test, and p-values were corrected for multiple testing using the Storey method ^8,26,27^. For bone analysis, protein groups with at least two unique peptides, q-value <0.05, and absolute log2(fold-change) >0.58 were considered to be significantly altered. Hippocampal analysis used a log2(fold-change) threshold of >0.2 (**Supplementary Tables 7-10**). Partial least squares-discriminant analysis of the proteomics data was performed using the package mixOmics in R (version 4.0.2; RStudio, version 1.3.1093)^6^.

### **Ingenuity Pathway Analysis**

Pathway analysis was completed on sets of differentially regulated proteins with the Ingenuity Pathway Analysis tool (QIAGEN, [Hilden, Germany](https://www.google.com/search?client=safari&sca_esv=02a881f72cac3b2d&rls=en&sxsrf=AE3TifNrSrfzWSR9UfMO2GmEMrzOJukxRA:1749594850946&q=Hilden&si=AMgyJEveiRpRWbYSNPkEPxCUbItHSvun4xkRgDDPLmrOjDx35LAse6695VLbg9RbUSQEb7JWAaEpHLEJ5pe_rrUX8JKlc4w6UWCJG8r4W8DTBU3aXKA6QelYMg3SSMxJvVRLBvOJ8paPvtTdsNVOey2HxBEf80Ev0YC5Zax7ktLzviIRENYIiShopI93OGG9rvz9XE375q2P&sa=X&ved=2ahUKEwilut3c9OeNAxWYIDQIHQsFJfUQmxN6BAg7EAI)). Pathway enrichment is scored by q-value, gene ratio, and z-score – a predictive score that measures whether a pathways is predicted to be up or down regulated.^28^ Enriched pathways with a significant corrected p-value (p<0.05) are shown in figures, with redundant pathways containing the same proteins represented by the pathway with the lowest p-value.

### **Weighted co-expression network analysis (WGCNA)**

### Protein co-expression network analysis was done utilizing log normalized protein abundance with the WGCNA R package^29^ as described^30^. Composite-tissue analysis was completed treating combined datasets as a single data set for cluster identification (**Supplemental Table 11**). Tissue-annotated cluster proteins were used to complete tissue specific pathway analysis as described above utilizing IPA.

**Data Availability**

RNAseq data generated from mouse cortical bone enriched in osteocytes are available in the National Center for Biotechnology Information (NCBI) Sequence Read Archive (SRA) under BioProject accession number PRJNA1314782.

Raw data and complete MS data sets have been uploaded to the Mass Spectrometry Interactive Virtual Environment (MassIVE) repository, developed by the Center for Computational Mass Spectrometry at the University of California San Diego, and can be downloaded using the following link: <ftp://MSV000099018@massive-ftp.ucsd.edu> (MassIVE ID number: **MSV000099018**; Proteome Xchange ID: PXD068037). **[Note to reviewers]:** To access data prior during revision (before public release) please utilize the following information: Username: **MSV000099018_reviewer***,* Password: **winter**.
